# Widespread horizontal gene transfer between plants and their microbiota

**DOI:** 10.1101/2022.08.25.505314

**Authors:** Shelly Haimlich, Yulia Fridman, Hitaishi Khandal, Sigal Savaldi-Goldstein, Asaf Levy

## Abstract

Plants host a large array of commensal bacteria that interact with the host. The growth of both bacteria and plants is often dependent on nutrients derived from the cognate partners, and the bacteria fine-tune host immunity against pathogens. This ancient interaction is common in all studied land plants and is critical for proper plant health and development. We hypothesized that the spatial vicinity and the long-term relationships between plants and their microbiota may promote or even depend on cross-kingdom horizontal gene transfer (HGT), a phenomenon that is relatively rare in nature. To test this hypothesis we analyzed the *Arabidopsis thaliana* genome and its extensively sequenced microbiome to detect events of horizontal transfer of full length genes that are absent from non-plant associated bacteria. Interestingly, we detected 180 unique genes that were horizontally transferred between plants and their microbiota. Genes transferred from plants to their microbiota are enriched in secreted proteins that metabolize carbohydrates, whereas microbes transferred to plants genes that are enriched in redox homeostasis functions. To validate our approach, we tested if a bacterial gene is functionally similar to its Arabidopsis homologue *in planta*. The Arabidopsis *DET2* gene is essential for biosynthesis of the brassinosteroid phytohormones and loss-of-function of the gene leads to dwarfism. We found that expression of the *DET2* homologue from *Leifsonia* bacteria of the Actinobacteria phylum in the *Arabidopsis det2* background complements the mutant, and leads to normal plant growth. Together, these data suggest that cross-kingdom horizontal gene transfer events shape the interactions between plants and their microbiome.

**Significance statement:** What are the genes that shape host-microbe interactions and what are their origins are fundamental questions in molecular ecology and evolution. We explored the evolutionary mechanisms that formed Arabidopsis-microbiota interactions, as a model for host-microbe interactions. We found prevalent horizontal gene transfer, affecting 180 genes, that occurred between plants and their commensal microbiota. We propose that these genes participate in molecular mimicry between the host and its microbiome. Bacteria acquired from plants genes that primarily encode for secreted proteins that metabolize carbohydrates, thereby enabling bacteria to grow on plant-derived sugars. Additionally, we demonstrate how a bacterial gene that mimics a plant hormone biosynthesis gene can replace the plant gene function. Our results suggest that horizontal gene transfer between hosts and their microbiota is a significant and active evolutionary mechanism that contributed new traits to plants and their commensal microbiota.

## Introduction

Plants form intimate associations with microbes, collectively called the plant microbiota. Microbes mostly have commensal lifestyles with the plant. However, the microbial molecular mechanisms used to interface with the host molecular network are mostly elusive. Successful isolation and subsequent genome sequencing of hundreds of bacterial strains that are associated with diverse plant species and tissues enabled the elucidation of some of the microbial genes that are responsible for plant adaptation (1–6). Transcriptome and proteome studies detected which bacterial genes are active *in planta (7, 8)*. As part of the interaction between plants and their microbiota different molecules are being exchanged, including simple carbohydrates, organic acids, signaling molecules, antimicrobials, and bacterial effector proteins that change plant physiology (9–16). Some of the exchanged molecules are used in molecular mimicry in which the microbes mimic the host or vice versa, to provide the acceptor with the donor’s biological functions, such as new metabolic capacity, and quick adaptation to the shared environment. Molecular mimicry was described extensively in plant pathogens. The *Pseudomonas syringae* pathogen uses its phytotoxin coronatine to structurally mimic the plant hormone jasmonoyl isoleucine (JA-Ile) and thereby to manipulate the host jasmonic acid signaling (17, 18). *P. syringae* also acquired a eukaryotic E3 ubiquitin ligase domain as part of an effector protein AvrPtoB that degrades a host kinase, leading to host disease susceptibility (19). Another pathogen, *Xanthomonas oryzae*, mimics the plant growth-stimulating PSY peptide to facilitate its infection (20). The commensal *Bacillus subtilis* encodes a remote homologue of plant expansin proteins. The bacterial expansin promotes plant cell wall extension and is critical for root colonization (21). Molecular mimicry can occur through cross-kingdom horizontal gene transfer (HGT) events (22–24). HGT can potentially occur between the plant host and its microbiome. The close proximity between the organisms, the selective pressure caused by immense microbial competition in the rhizosphere and by the plant immunity, and the inherent ability of microbes to integrate foreign DNA may guide DNA transfer from plants into their microbiome (25). Gene transfer events from bacteria into plant germ cells have an unclear mechanism. However, such events were reported in the past and were described as mostly ancient HGT events from bacteria into the ancestors of land plants (26–29). Of course, *Agrobacterium* transfers genes into somatic plant cells but these genes do not pass to the plant seeds.

We hypothesized that the genomes of plants and their direct root and shoot microbiome could uncover the extent and nature of cross-kingdom HGT events between plants and bacteria. In the current study we performed a systematic search for potential cross-kingdom HGT events between the model plant *Arabidopsis thaliana* and its extensively isolated and sequenced microbiome. Using a phylogenetic analysis of hundreds of proteins from plants, plant-associated bacteria, and several control groups we could detect 180 horizontal gene transfer events and determine their directionality and their taxonomic and functional biases. Moreover, we showed that a gene that was transferred from plants to plant-associated bacteria still maintains its *in planta* function despite its divergence from the plant homologue through mutations. Our data suggests that cross-kingdom HGT events are relatively frequent, and shaped the genomes of plants and their microbiota by adding adaptive traits to the recipient organisms.

## Results

### Identification of genes that share high amino acid sequence similarity between *Arabidopsis thaliana* and its microbiome

To quantify the extent of molecular mimicry events between plants and microbiome we focused on the plant *Arabidopsis thaliana* which serves as a model for plant-microbe interactions (30, 31). The microbiota of Arabidopsis were extensively isolated from roots and shoots by different groups and their genomes were previously sequenced (1, 2, 4). We compared the genes of *A. thaliana* against the genes of 582 fully sequenced bacteria that were isolated from *Arabidopsis thaliana* (Supplementary Table 1) Our current analysis includes commensal Arabidopsis-associated bacteria which were isolated from healthy plants, whereas most molecular mimicry events were previously studied in the context of plant pathogens such as the *Xanthomonas* genus and *Pseudomonas syringae* species (32). We used BLASTP program to detect similarity between the set of proteins encoded by Arabidopsis and its bacteria (Figure 1A). We focused on similarity between full-length protein sequences, as these proteins are relatively poorly studied in comparison to protein domains involved in molecular mimicry. Protein domains are extensively studied in the context of effectors of pathogenic bacteria (33, 34). We used only BLASTP hits of at least 35% amino acid sequence identity across at least 80% of both the Arabidopsis and the bacterial proteins. Next, we filtered out what we termed ‘trivial hits’ which are protein sequences shared by bacteria and plant organelles, mitochondria and chloroplast, which themselves originate in bacteria (35), and their proteome may maintain high similarity to their original bacterial proteome. This analysis resulted with a list of 767 *Arabidopsis* proteins that are mapped to 60,850 bacterial proteins and likely play similar functions (Supplementary Table 2). One concern is that protein sequence similarity-based search will result in detection of Arabidopsis or bacterial DNA contaminants that were mistakenly assembled in bacterial or Arabidopsis genomes, respectively. However, we did not identify bacterial-Arabidopsis homologous protein pairs with amino acid identity above 76%, rejecting the possibility of hypothetical DNA contamination which would lead to highly similar protein sequences.

**Figure 1.**
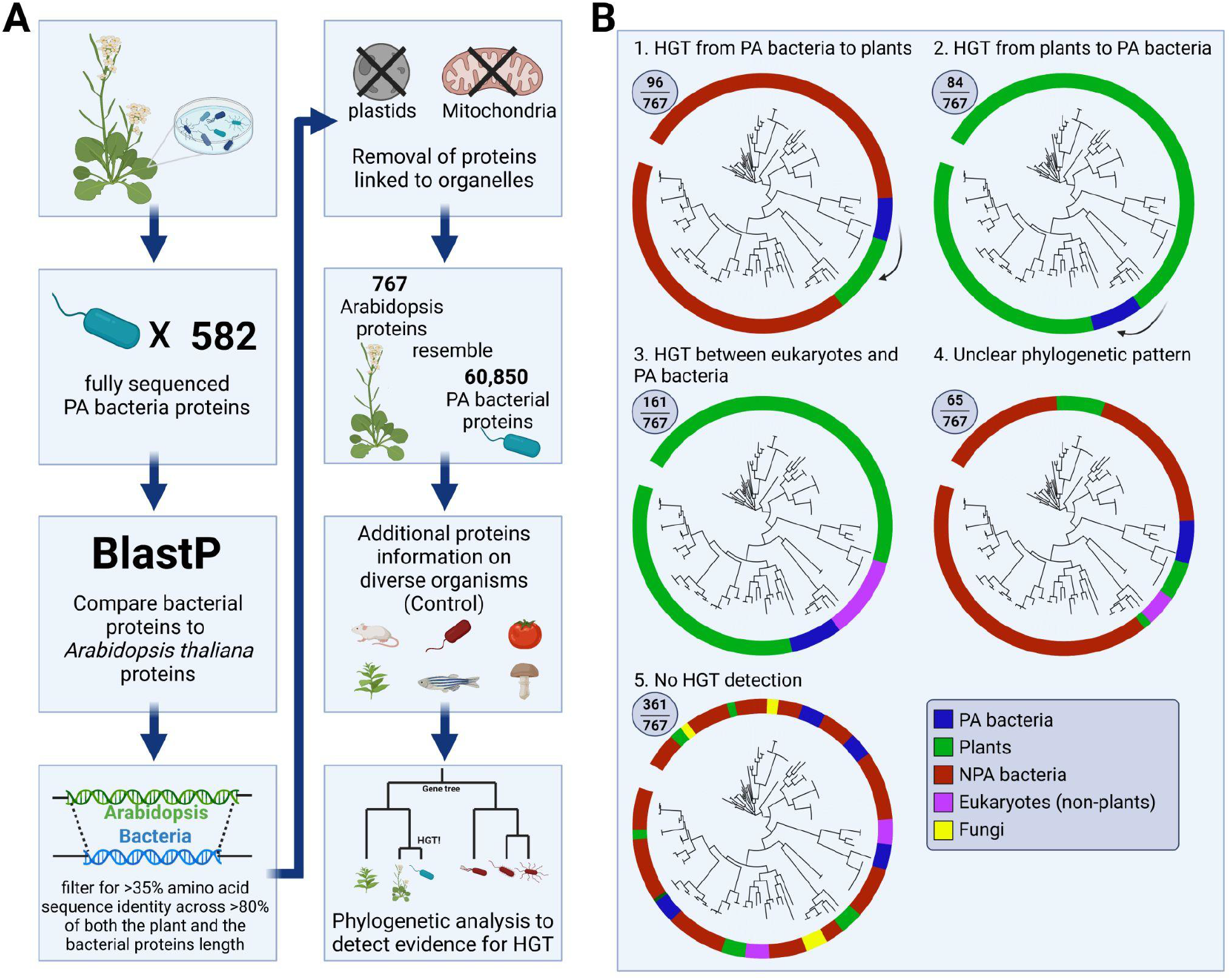
Analysis used to detect cross-kingdom HGT events between plants and bacteria. **A**. An outline of the bioinformatic analysis we performed to detect molecular mimicry events and cross-kingdom HGT events between plants and their microbiota. **B**. Classification of the molecular mimicry events into different classes based on their evolutionary path. PA - plant-associated, NPA - non-plant associated.

### One hundred eighty genes demonstrate cross-kingdom horizontal transfer from plants to their microbiota or vice versa

The proteins we identified as sharing high sequence similarity between plants and their microbiota can be the result of various evolutionary scenarios. For example, the genes can be ancient and conserved between eukaryotes and prokaryotes. We specifically searched for phylogenetic evidence supportive of direct transfer across plants and their microbiota. To this end we compiled a dataset that contained 1051 genomes (Supplementary Table 1), including 1023 bacteria from Actinobacteria, Proteobacteria, Firmicutes, and Bacteroidetes phyla, ten monocot and dicot plants, eight fungi, two archaea and eight additional eukaryotes (animals, parasites, and a mold). Importantly, the bacteria were divided into plant-associated (“PA”, n=582, mostly isolated from *Arabidopsis*) and non-plant associated (“NPA”, n=441, isolated from elsewhere) based on their original isolation site, which we previously manually curated (2). Gene transfer events between plants and PA bacteria enable identification of the habitat where the transfer event occurred and to assign a likely adaptive role in the plant environment to the transferred gene.

For each of the 767 Arabidopsis genes involved in predicted molecular mimicry we constructed a gene tree based on multiple sequence alignment of all gene hits within our genome dataset (Supplementary Dataset 1). We defined an event of cross-kingdom horizontal gene transfer from plants to their microbiota or vice versa when a subset of organisms from both groups shared the same branch in the gene tree demonstrating an inconsistency between the genome tree and the gene tree (Figure 1B). Specifically we verified that the branch that is composed of homologues from plants and PA bacteria did not include homologues from animals or NPA bacteria. We hypothesize that having a gene shared only between plants and their microbiota increases the likelihood that the gene was directly transferred and has been maintained in the acceptor genome despite the long phylogenetic distance due to its function in the plant environment. However, we cannot reject a possible scenario of massive gene loss in the non-plant environment.

We defined the donor doman (e.g. bacteria) based on the organisms in the closest sister clades of the branch that contains plants and PA bacteria. Overall, we identified 84 genes that were likely transferred from plants to their bacteria and 96 genes that were likely transferred from PA bacteria to their plant hosts (Supplementary Table 3, Supplementary Figures 1-6). In addition, we identified 161 genes that were horizontally transferred between PA bacteria and the eukaryotic domain; in these cases, it is not possible to determine whether the transfer occurred between plants and PA bacteria or another eukaryotic group and PA bacteria (Supplementary Table 3, Supplementary Figures 7-9). Finally, we classified two more groups; The first contains 65 genes whose pattern of inheritance is unclear and can be interpreted in more than one way, and the second group includes 361 genes in which no HGT patterns were observed (Supplementary Table 3, Supplementary Figures 10-15).

### Genes that were transferred from plants to bacteria and from bacteria to plants are enriched in carbohydrate metabolism and redox homeostasis functions, respectively

We performed a functional enrichment analysis of the genes that have been horizontally transferred (Materials and Methods). The genes transferred from plants to PA bacteria are enriched in genes encoding secreted proteins (Figure 2A), including proteins with Gene Onthology (GO) terms “signal”, “secreted”, and “extracellular region”. These genes are also enriched in enzymes that perform carbohydrate catabolism with pectin esterase or glycosidase activity and they likely target the plant cell wall or exploit carbohydrates from the root exudates (36). These genes include, for example, genes encoding chitinases (CHI, AT2G43570), glycosyl hydrolases such as an endo beta mannanase (XCD1, AT3G10890), and pectin lyases (PME5, AT5G47500). The chitinase (CHI) gene also serves as a defense gene that is induced in plants during Systemic Acquired Resistance (37, 38) and its transfer to bacteria may be used to manipulate plant defense response or to directly attack fungi in the plant environment. In contrast, the genes that were transferred from PA bacteria to plants are mostly found in all dicots and monocots that we analyzed (69/96 genes, 72%), suggesting that these are ancient HGT events. However, 10 Arabidopsis genes that originate in PA bacteria are found only in dicots based on our analysis (Supplementary Table 3). These genes include, for example, AT1G24290 which encodes for a AAA-type ATPase family protein. Previous works also suggested that genes were transferred from bacteria to the ancestors of land plants (27, 29). We detected within this list many of the auxin biosynthesis YUCCA genes (YUC1, YUC3, YUC5, YUC6, YUC7, YUC8, YUC9, YUC11) that were previously suggested to be acquired from bacteria, as a consequence of plant interaction with microbes (27, 39). In contrast to genes transferred from plants to bacteria, we identified that genes transferred in the other direction are enriched in genes that encode cytoplasmic proteins (Figure 2B). In addition to auxin biosynthesis related functions, genes transferred to plants are enriched in nucleoside metabolic processes, such as uridine-ribohydrolase 1 (AT2G36310). We analyzed the number of introns found in Arabidopsis genes transferred from PA bacteria assuming that they would contain less introns than the average plant gene. However, we found no statistical difference between the number of introns of horizontally transferred genes and all other Arabidopsis genes (Supplementary Figure 16). This result can further support the hypothesis that the gene transfer into plant is ancient enough so that these genes now resemble the rest of the plant genes.

**Figure 2:**
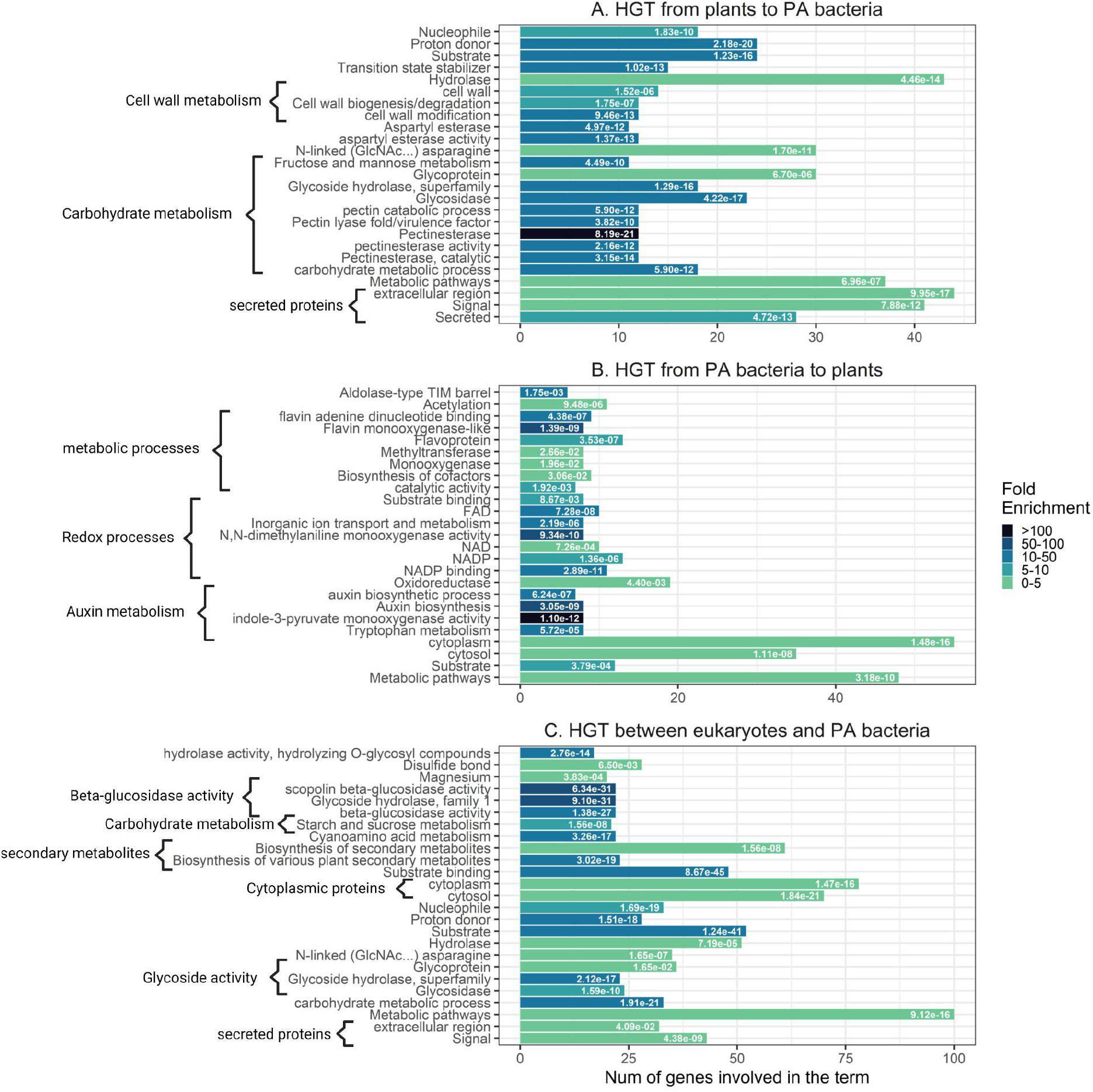
Gene Ontology (GO) enrichment analysis of the different groups that have signatures of HGT. GO enrichment analysis using DAVID. The 25 most significantly (P<0.05, after false discovery rate correction) enriched GO terms. The fold wnrichment ia shown in green to black color scale and the q-values are written in white on the bars **A**. GO functional analysis of Arabidopsis genes (n=84) with an HGT pattern from plants to PA bacteria. **B**. GO functional analysis of Arabidopsis genes (n=96) with an HGT pattern from PA bacteria to plants. **C**. GO functional analysis of Arabidopsis genes (n=161) with an HGT pattern between PA bacteria and the eukaryotic domain.

A third group of genes includes more complex HGT scenarios, from plants or other eukaryotes to PA bacteria. This group is enriched in genes that encode cytoplasmic proteins and proteins with glycoside hydrolase or beta-glucosidase activity (Figure 2C).

### Taxonomic patterns of the molecular mimicry events

We next asked if specific microbes tend to be gene acceptors or gene donors. We performed an enrichment analysis considering the taxonomy of bacteria that underwent HGT using the entire bacterial genome set as controls (Materials and Methods). First, we analyzed HGT events from plants to PA bacteria at the phylum level (Figure 3A). Actinobacteria and Proteobacteria were depleted for being gene acceptors, whereas Bacteroidetes were enriched as gene acceptors. Interestingly, in most cases the enriched taxonomic group was”unknown” as microbes from more than one group are located at the branch that received the gene from plants. Using the current taxonomic information we cannot tell if this HGT pattern is the result of gene transfer to one microbial phylum followed by an additional transfer to another microbial phylum, or independent gene transfer event to multiple phyla. When increasing the taxonomic resolution and looking at the order taxonomic rank the pattern of unknown gene acceptor persisted as being enriched (Figure 3A). An example can be seen for gene AT1G54310.2 (Figure 3B), which encodes an S-adenosyl-L-methionine-dependent methyltransferases superfamily protein. The gene is conserved in monocots, dicots, and mosses and was likely horizontally transferred from moss (*Physcomitrium patens*) into a limited group of Bacteroidetes (*Pedobacter*) and Proteobacteria (*Rhodanobacter*) species. For three orders we observed significantly less HGT events as expected by the number of sequenced genomes from these orders: Rhizobiales and Hyphomicrobiales (Proteobacteria) and Micrococcales (Actinobacteria). Next, we analyzed the biases in bacterial taxonomy in HGT events from PA bacteria into plants. We detected an enrichment of events where the donor was of unknown taxonomy (namely, multiple phyla can serve as the donors) and a depletion of Proteobacteria and Actinobacteria serving as gene donors. These biases were very similar to the scenario of PA bacteria serving as gene acceptors. Since the transfer from PA bacteria to plants was relatively ancient we could not perform this analysis at taxonomic levels below phylum. To summarize this analysis, in most cases the taxonomy of the bacterial donor or acceptor is unknown, and Actinobacteria and Proteobacteria relatively rarely donate or accept genes from plants. This trend includes members of the Rhizobiales order which serve as common endophytes but in comparison to other groups Rhizobiales did not accept many genes from plants.

**Figure 3.**
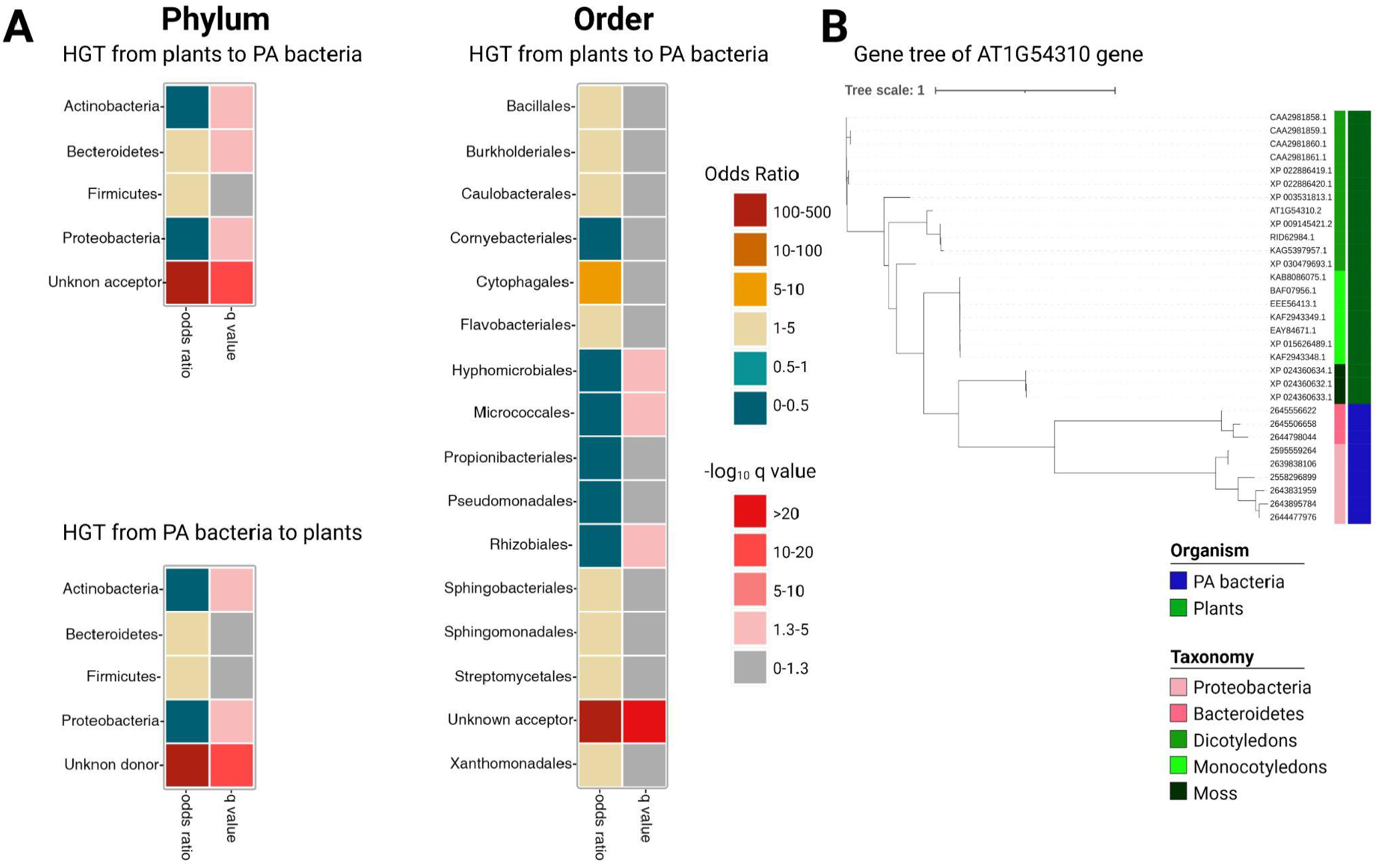
**A**. A Fisher exact test was used to examine bacterial taxonomic groups that tend to accept/donor genes. The Odds Ratio represents the enrichment or depletion magnitude, with orange/red colors representing enrichment, and blue colors representing depletion. The negative log10 of the q-values, shown in gray to red colors, are corrected for multiple hypothesis testing, while gray color represents non significant values. **B**. An example of a plant gene (AT1G54310) transferred from plants to PA bacteria from two phyla (“Unknown acceptor**”)**.

### Signatures of recent horizontal gene transfers into bacterial genomes can be detected

In several cases we identified HGT events from plants into plant-associated bacteria that likely occurred relatively recently. These events were characterized by insertion into a single plant-associated bacterial genus. In addition, through inspection of the genomic neighborhood of the acquired gene we observed that the gene had a patchy presence/absence pattern between members of the same genus and was located in a relatively variable genomic region compared to its flanking regions. One example is the bacterial homologue of the plant-specific *CHI* gene (AT2G43570), encoding a putative basic chitinase. *CHI* gene is a defense gene that is strongly upregulated in plants in response to butterfly oviposition (40). The *CHI* gene is present in seven genomes of PA *Streptomyces* genomes but is absent in other PA *Streptomyces*, suggestive of a recent gene acquisition/loss event (Fig. 4A). Comparison of the predicted protein structures of the plant and bacterial CHI homologues demonstrate a striking similarity with Root-Mean-Square-Deviation (RMSD)=0.791 (Fig. 4C). The N-terminus presented the largest difference between the two structures.

**Figure 4.**
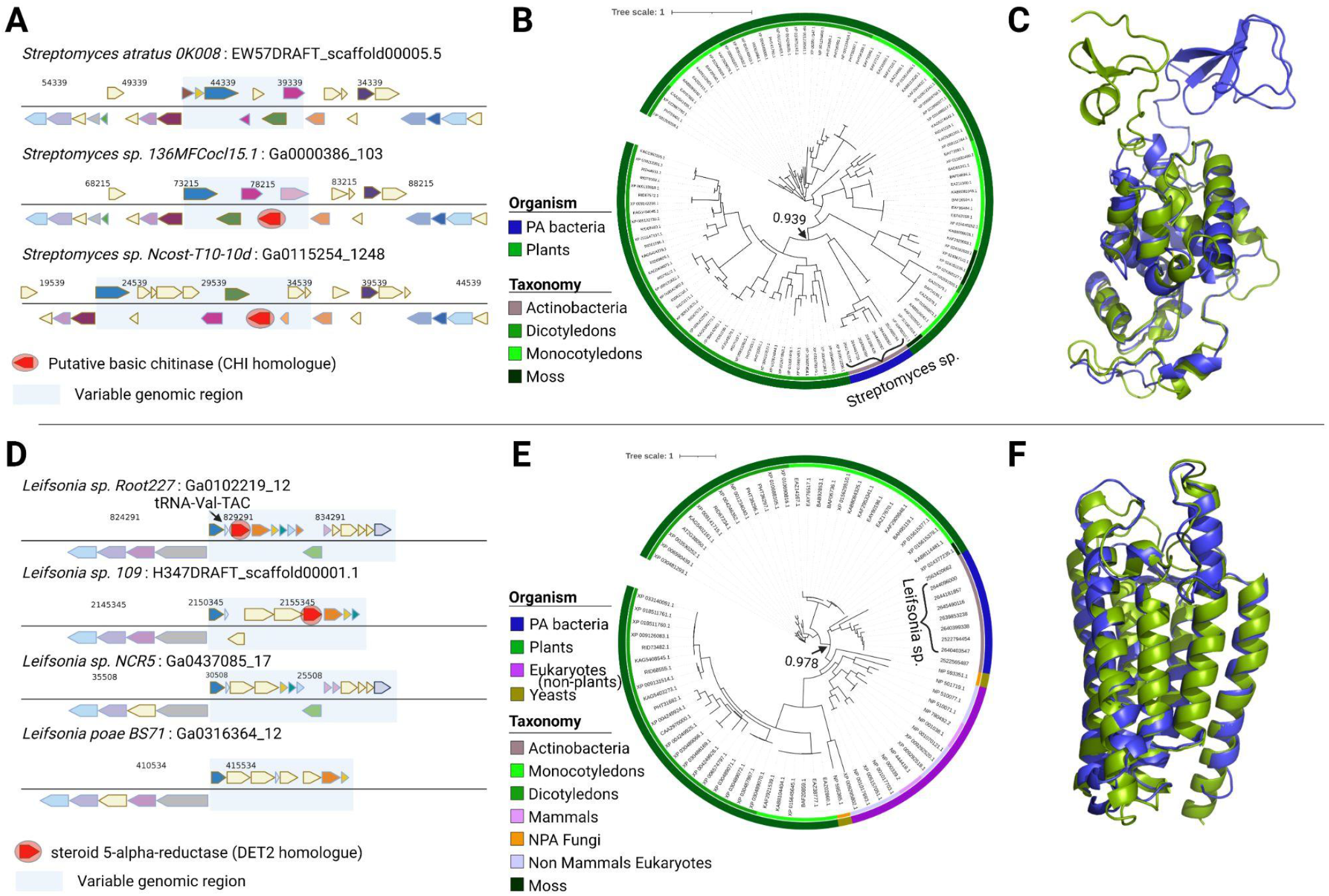
Examples of likely recent HGT from plants or other eukaryotes to PA bacteria. **A**. Pattern of presence/absence of the CHI homologue gene (marked with a red circle) in PA *Streptomyces* genomes. **B**. A phylogenetic tree that presents CHI protein found mainly in plants and also found in a small group of PA *Streptomyces*. The bootstrap value of the clade that is shared by plants and their bacteria is 0.939 (marked with an arrow). **C**. Structure comparison of CHI plant protein (green) with bacterial protein (blue), Root-Mean-Square Deviation= 0.791. The top part represents the N-terminus. **D**. Pattern of presence/absence of the *DET2* homologue gene (marked with a red circle) in *Leifsonia* genomes. **E**. A phylogenetic tree that presents DET2 proteins found mainly in the eukaryotic domain and in a small group of *Leifsonia* bacteria. The bootstrap value of the clade that is shared by plants, their bacteria, and other eukaryotic organisms is 0.978 (marked with an arrow). **F**. Structure comparison of DET2 plant protein (green) with bacterial protein (blue), Root-Mean-Square Deviation = 0.727.

Another interesting example is of the bacterial homologue of *DET2*, a steroid-5-alpha-reductase which is one of the key genes in the brassinosteroid biosynthetic pathway (41). The bacterial *DET2* is nearly specific to plant-associated *Leifsonia* genus from Actinobacteria phylum (Figure 4E) and it is the only gene from the brassinosteroid biosynthetic pathway that we detected in bacteria. The *Leifsonia DET2* branch diverged from plants and is located next to a non-plant eukaryotic branch (Figure 4E). Therefore, it is difficult to determine if the gene was acquired directly from plants or from other eukaryotes. The gene is located in a variable genomic region downstream to a tRNA gene (Figure. 4D) which may mediate integration into the locus of foreign DNA (42). Although the plant and bacterial DET2 homologues share less than 50% sequence identity their predicted structures are strikingly similar with Root-Mean-Square-Deviation (RMSD)=0.727, suggestive of a similar biochemical function (Figure 4F).

### Horizontally transferred bacterial gene can functionally replace its homologous plant gene

We tested if the bacterial genes we identified as being horizontally transferred from plants can replace their homologous plant genes. As a proof-of-concept we selected *DET2* (Figure 4D-F). The Arabidopsis DET2 protein is 46% identical to the *Leifsonia* Det2 homologue. Interestingly, this level of sequence similarity is shared between the Arabidopsis DET2 and its homologues from rice and barley. Importantly, the proteins from other eukaryotic plant pathogens such as *Phytophthora* and *Pythium* share weaker identity (maximum 41% identity) to the Arabidopsis protein than the Leifsonia-Arabidopsis DET2 similarity. The *det2* mutant has a severe dwarf phenotype including a wider root meristem with altered cell wall orientation, typical to the BR deficient mutants (Figure 5A-C) (43, 44). We transformed this mutant background with Leifsonia Det2 (lfDET2) fused to a fluorescent protein (lfDET2-NG), driven by the constitutive 35S or the Arabidopsis DET2 promoters and observed a rescue of these BR phenotypic defects (Figure 5A-C). The rescued root length remained shorter than wild type, similar to an equivalent transformation with the Arabidopsis DET2 (*atDET2*) (45). In agreement with the atDET2-like functionality *in planta*, lfDET2 localized to the endoplasmic reticulum, similar to *atDET2* (Figure 5D-F). To conclude, lfDET2 functionally replaces its Arabidopsis homologue.

**Figure 5.**
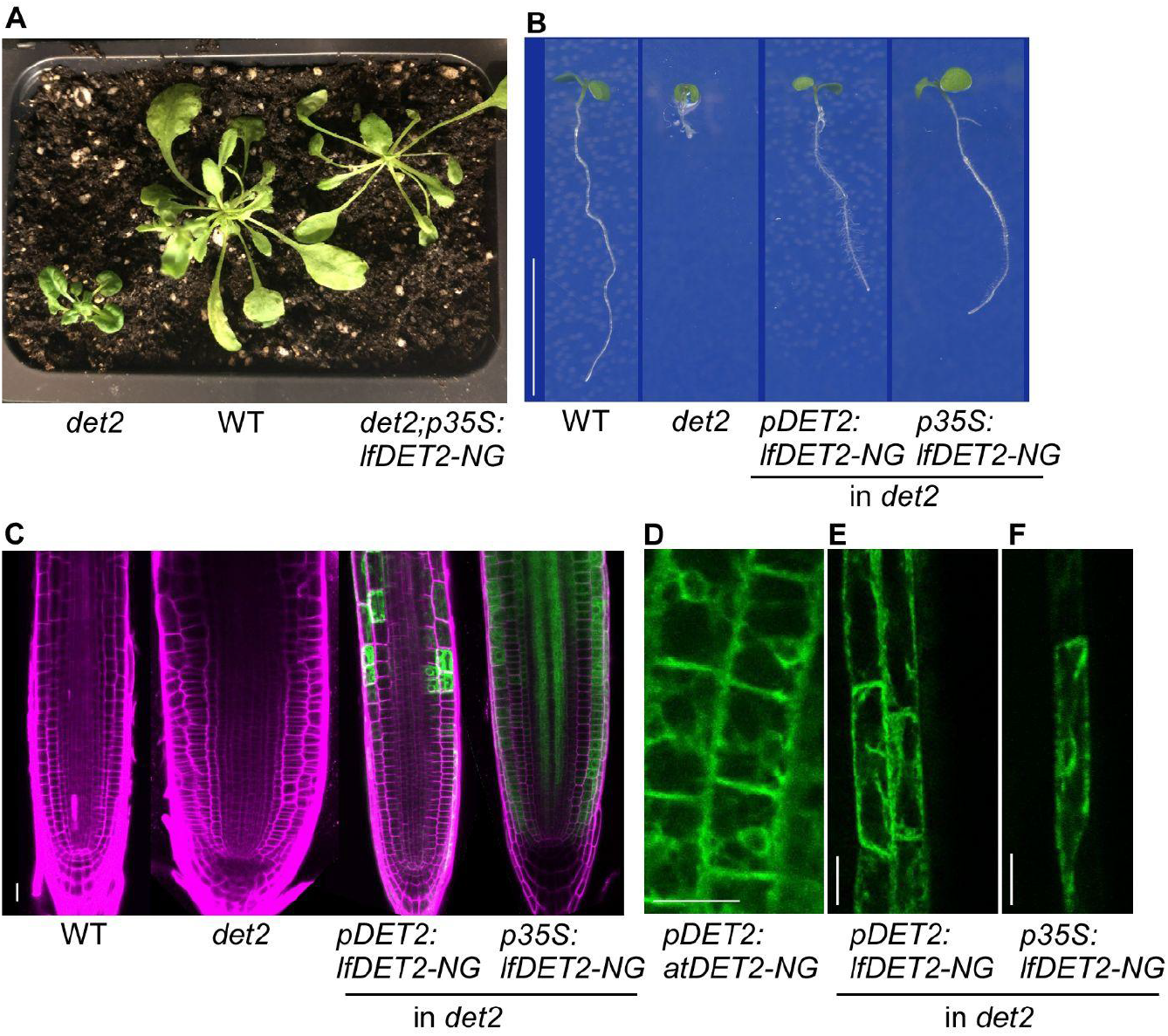
Complementation of *det2* Arabidopsis by a *det2* bacterial homologue reveals functional similarity between bacterial and Arabidopsis homologous genes. A comparison between wild type (WT), *det2*, and transgenic *det2* lines harboring *pDET2:lfDET2-NG* and *p35S:lfDET2-NG* (*det2* expressed from two promoters). **A**. Adult developmental stage of *det2*, WT and *p35S:lfDET2-NG*. Note the WT-like phenotype of the rescued *det2*. **B**. 7-day-old seedlings of WT, *det2, pDET2:lfDET2-NG* and *p35S:lfDET2-NG*. Scale bar = 1 cm **C**. Root meristem of lines as in **A**. Note the wide and aberrant morphology of the *det2* meristem and its rescue by lfDET. mNeonGreen(NG) is shown in green and propidium iodide (PI) that marks cell borders in magenta. **D-F**. Subcellular localization of atDET2 and lfDET2 in epidermal root cells. Note their similar localization in the endoplasmic reticulum. Scale bars = 20 um.

## Discussion

In this work we looked closely on the effect of HGT on the evolution of plants, with a focus on *Arabidopsis thaliana* and its extensively sampled microbiome. The effect of HGT on plant evolution was described in several works in recent years. Analysis of the moss *Physcomitrella patens* identified 44 families of nuclear genes that were acquired from bacteria in comparison to only 10 gene families that were acquired from fungi, and only one family is from archaea and another was acquired from viruses (27). These findings include two gene families that are involved in auxin biosynthesis. Green plants acquired from bacteria genes related to biosynthetic and metabolic pathways, adaptation, exaptation, and stress response (46). These include, for example, genes involved in xylan degradation, plant vascular system development, and stress response to cold and cadmium. Genome analysis of subaerial Zygnematophyceae algae concluded that gene families that increase resistance to biotic and abiotic stresses in land plants, in particular desiccation through abscisic acid (ABA), were acquired by HGT from soil bacteria (47). A recent work analyzed HGT patterns in 12 representative plant species (48). The authors identified two major HGT episodes in plant evolution, each of which contributed more than 100 gene families. The first occurred during early evolution of streptophytes and the second at the origin of land plants. Most of the contributing organisms were described as bacteria, with some contribution from fungi. These results also support our current results that nearly all transfers into Arabidopsis are ancient and are shared by monocots and dicots. We could not detect a bacterial gene that was transferred directly to Brassicaceae and is absent from other dicots.

Previous works could not specifically determine which microbes (at a low taxonomic rank) were gene donors or acceptors. Therefore, we thought that the microbiome of a plant is the natural partner for HGT. We focused on bacteria and not on fungi and archaea, because there is a high number of sequenced Arabidopsis-associated bacteria, and because previous works showed that they are the most common gene donors to plants. Another addition to our approach is the separation between bacteria according to their isolation sites to plant-associated (PA) and non-plant associated (NPA) with the latter serving as controls. This distinction allowed us to suggest the path of the HGT to occur via the plant microbiome. In addition, in the case of gene transfer to PA bacteria it allowed us to suggest that the gene was maintained following transfer to increase bacterial fitness in the plant environment. Our observed enrichment of carbohydrate catabolism-associated gene transfer, specifically of plant sugars such as pectin and xylose, into PA bacteria supports the hypothesis that the new bacterial genes play an adaptive role. These genes, including the genes for pectate lyases, glycosyl hydrolases, sugar isomerase, and arabinanase, increased plant-dependent bacterial growth and likely the ability to colonize plant tissues by breaking down the plant cell wall. Pectate lyases enzymes were also shown to be critical for endophytic Arabidopsis root colonization by fungi, and they reduced plant performance (49).

Interestingly, in a previous study we showed that genomes of PA bacteria have a higher number of carbohydrate metabolism genes than the genomes of the NPA group from the same taxon (2). This trend was reproducible when we examined groups of bacteria from four different phyla. In the current work we propose a model that actually some of these genes were transferred directly from plants to their microbiome. For example, Cluster of Orthologous Genes (COG) 4677 describes a large group of Pectin methylesterase and related acyl-CoA thioesterases genes that alter cell wall integrity. We identified COG4677 as being in higher presence in PA bacteria than in NPA bacteria from the Bacteroidetes phylum, Burkholderiales and Bacillales orders, and Xanthomonadaceae family (2). In the current work we saw that these genes were transferred from plants to PA bacteria, including the pectinesterases PME5, PME31, PME44, QRT1 and others, that have homologous bacterial proteins of 35%-40% sequence identity.

Several phytohormone pathways have previously been described as transferred between plants and associated microbes. Agrobacterium naturally transfers its auxin biosynthesis genes into its host plant (50). It was suggested that the YUC genes that are involved in Auxin biosynthesis were transferred from bacteria to the most recent common ancestor of land plants (27). Agrobacterium also encodes and transfers into plants genes required for production of cytokinin (51). On the other hand, as part of the interaction with plants, rhizobia evolved an independent pathway for gibberellin production (52). In the current work we showed that *DET2*, from the brassinosteroid biosynthetic pathway may have transferred to plant-associated bacteria, mostly to the *Leifsonia* genus. We showed that the gene can replace its plant homologue in brassinosteroid production but the role of this gene in bacteria remains unknown. We performed Arabidopsis root colonization experiments with *Leifsonia* strains that are naturally positive or negative for *det2* and could not detect a phenotypic effect that correlates with the gene presence. Thus, the significance of this HGT remains unclear. Together, our results suggest that HGT shaped the genomes of both plants and associated microbiota and that specific gene functions were acquired by bacteria, likely to utilize the unique carbohydrate-rich environment that plants produce due to photosynthesis followed by root exudation into the surrounding soil.

## Materials and Methods

### Data sources and genome screening

Full genomic sequence of Arabidopsis thaliana was downloaded from TAIR website (https://www.arabidopsis.org/). Additionally, a collection of 582 microbiota was prepared. These bacteria from the roots and shoots of Arabidopsis thaliana were extensively isolated by different groups, and their genomes were previously sequenced. Using BLASTP (53) version 2.8.1+ (standard settings), a comparison was made between the 582 bacterial and Arabidopsis proteins. The hits were filtered of at least 35% amino acid sequence identity across more than 80% protein length of the plant and the bacterial proteins. Protein sequences shared by bacteria and plant organelles, which themselves originate in bacteria, were filtered out. Information on plant organelles origin is taken from ATH_GO_GOSLIM.txt file on the TAIR website (https://www.arabidopsis.org/download/GO and PO Annotations/Gene Ontology Annotations /ATH_GO_GOSLIM.txt.gz). At the end of this analysis, we identified 767 Arabidopsis proteins that were mapped to 60,850 bacterial proteins (Supplementary Table 2).

### Phylogenetic comparative methods to find HGT

A dataset that contains fully sequenced diverse organisms was created. The dataset contained 1051 genomes, including 1023 bacteria (582 PA bacteria, 441 NPA bacteria), 10 monocot and dicot plants, 8 fungi, 2 archaea, and 8 additional eukaryotes (Supplementary Table 1). Automatically, multiple sequence alignment was performed using Clustal Omega (54) version 1.2.4 using standard settings, and each of the 767 phylogenetic trees was constructed using FastTree (55) version 2.1.11 SSE3 using standard settings (Supplementary Dataset 1). Display, annotation, and management of phylogenetic trees were performed with Interactive Tree Of Life (56) (ITOL version 6.5). Each of the phylogenetic trees was examined manually, and all the trees were divided into five categories according to the inheritance pattern observed in the tree: (1) HGT from PA bacteria to plants (2) HGT from plants to PA bacteria (3) HGT between eukaryotes and PA bacteria (4) Unclear phylogenetic pattern (5) no HGT detection (Supplementary Table 3).

### Gene Ontology enrichment analysis

Functional information about the gene was performed using a function annotation test (GO analysis) using DAVID (57) version 6.8 on the Arabidopsis genes in three different groups: HGT from PA bacteria to plants, HGT from plants to PA bacteria and HGT between eukaryotes and PA bacteria.

### Enrichment analysis

To test which bacteria tend to accept or donor genes, each phylogenetic tree was examined manually to find the PA bacteria taxonomic group in the branch shared with plants. All taxonomic levels were examined to find the lowest taxonomic level common to all PA bacteria in the same plant branch. If two different Phyla were on the same branch, the acceptors or donors were determined to be “Unknown”. Fisher exact test was performed using python package scipy.stats, and performing an FDR correction to calculate the q-values.

### Genomic neighborhood, structure prediction and structure comparison

A pattern of genes present/absent was created by using the IMG website (58), using the option “Show neighborhood regions with the same top COG hit (via top homolog).” The structure prediction of the Arabidopsis proteins was downloaded from UniProt website (https://www.uniprot.org/), and the prediction structure of the bacterial protein was created using AlphaFold2 (59, 60) The structures comparison was created using PyMOL (52) version 2.4.1

### Graphs and Figures

Various R packages are used to create graphs: ggplot2, reshape2, dplyr, tidyverse and RColorBrewer.

BioRender (https://biorender.com/) was used to create figures 1, 3 and 4.

### Growth conditions, molecular cloning and transformation

For overexpression of *lfDET2*, the bacterial gene sequence underwent codon optimization for *Arabidopsis thaliana*. The constructs were generated using the Golden Gate MoClo Plant Tool Kit (61). For *DET2* promoter (*pDET2*), 550 bp fragment upstream to the first *DET2* ATG was used. *pDET2* and *p35S* in level 0 (in pICH41295) were then subcloned along with additional level 0 parts: lfDET2 (in pAGM1287), mNeonGreen (NG, in pAGM1301) and the *RBCS* terminator (in pICH41276) into level 1 (pICH47742). For overexpression of the Arabidopsis *DET2* (*atDET2*) a similar cloning procedure was used except that the *DET2* terminator was used. The constructed level 1 was then subcloned into a level 2 construct (pAGM4723), together with a level 1 kanamycin resistance gene (pICH47732). Plant transformation to wild type Col-0 and *det2* backgrounds was performed using the *Agrobacterium tumefaciens* (GV3101)-mediated floral dip transformation method. Transgenic lines were screened on selective 0.5 MS plates supplied with 50 mg/l kanamycin (Duchefa Biochemie). Homozygous lines were selected according to mendelian segregation of the selection marker. For each construct used, 2-3 independent transgenic lines were generated. Presented here are line 6 (*det2;pDET2:lfDET2*), line 3 (*det2;p35S:lfDET2*) and line 4 (*pDET2:atDET2*). Plant growth conditions were as described by Fridman et al. (62).Briefly, seeds were surface sterilized and germinated on one-half-strength (0.5) Murashige and Skoog (MS) medium supplemented with 0.8% plant agar, 0.46 g/l MES pH 5.8, 0.2% (w/v) sucrose. Plates with sterilized seeds were stratified at 4°C for 2 days in the dark before transfer to the growth chamber with 16 h light /8 h dark cycles, at 22°C. Irradiance conditions of ∼70 μmol m^-2^ s^-1^.

### Confocal microscopy

Confocal microscopy was performed using a Zeiss LSM 510 (Zeiss, Jena, Germany) confocal laser scanning microscope with a LD LCI Plan-Apochromat 25 × water immersion objective (NA-0.8), or LSM 710 (Zeiss, Jena, Germany) using a Plan-Apochromat 20x objective (NA-0.8). Roots were imaged in water, or with water supplemented with propidium iodide (PI, 10 μg/mL). The green fluorescent proteins NeonGreen (NG) and PI were excited by an argon laser (488 nm) and by DPSS laser (561 nm) respectively. For PI detection in LSM 710, solid state laser (543 nm) was used. In LSM 510, fluorescence emission signals for NG and for PI were collected by PMT detectors, with a band-pass filter (500-530 nm) and a long-pass filter (575 nm), respectively. In LSM 710, fluorescence emission signals for NG and for PI were collected by BIG detectors (GaAsP) with a band-pass filter (500-550 nm) and a band-pass filter (570-620 nm), respectively.

### lfDET2 sequence after codon optimization

ATGCCCGACGGTCCGTATCGCTGGTTCGTGTATGCCGAGATCGCCCTCGCGGTGG

TCACCTTCGTCGCTCTGTGCTTCGTGGTAGCGCCGTACGGACGGCACGGCCGCTC

CGGATGGGGGCCGACCGTGCCCGCGCGGGTCGGCTGGGTCGTGATGGAGAGTC

CAGCATCCATCGTCTTCCTGCTGTTCTACCTGCTCGGCGACCACCGGTTCGAGCTG

TGCCTCTGCTGTTCCTCGCGCTGTGGCAGCTCCACTACGTGCAGCGTGCCTTCG

TCTACCCGTTCCTGATGCGCACCGGGTCCAGGATGCCCGTGTCCGTCGTGGGGAT

GGCGATCCTGTTCAACCTGCTCAACGCGTGGGTGAATGCGCGGTGGATCTCGCAG

TACGGCCAGTACGCGAACAGCTGGCTCGCCGACCCTCGGTTCTGGATCGGCGTGG

TCGTGTTCATCGCCGGGTTCTCGCTCAACCTCGGTTCCGACCGCATCCTGCGCAG

ACTGCGGGGTGCGCGATCCGGCGGGTACAGCATCCCGCGCGGTGGCGGATACCG

CTGGGTGTCCAGCCCGAACTACCTGGGCGAGATGGTGGAGTGGACCGGCTGGGC

GATCGCGACCTGGTCGCTCGCCGGGCTGGCGTTCGCGCTGTACACGATCGCGAA

CCTCGCACCGCGGGCGATGGCGAACCACCGCTGGTACCTGGAGACGTTCGACGA

CTATCCGCCGGAGCGAAAAGCGATCATCCCCTATCTGCTCTGA

## Acknowledgments

The research was funded by a grant number 12-12-0002 from the Israeli Ministry of Agriculture awarded to AL and S.S.-G and by the Israel Science Foundation (no. 1725/18) to S.S.-G.. SH is supported by an excellence scholarship from the Faculty of Agriculture, Food, and Environment of the Hebrew University of Jerusalem. AL is also supported by the Alon Fellowship of the Israeli Council of Higher Education and the Israeli Science Foundation (grants 1535/20, 3300/20); Hebrew University - University of Illinois Urbana-Champaign seed grant; ICA in Israel. We thank the Life Sciences and Engineering Infrastructure Center (N. Dahan, Y. Lupu-Haber), the Russell Barrie Nanotechnology Institute at the Technion and O. Erlichman for helping with the last stages of the *det2* experiments. We thank Dr. Omri Finkel, Prof. Uri Gophna, and Prof. Itay Mayrose for their useful insights about the research.

## Supplementary Information

**Supplemental Figure 1.**
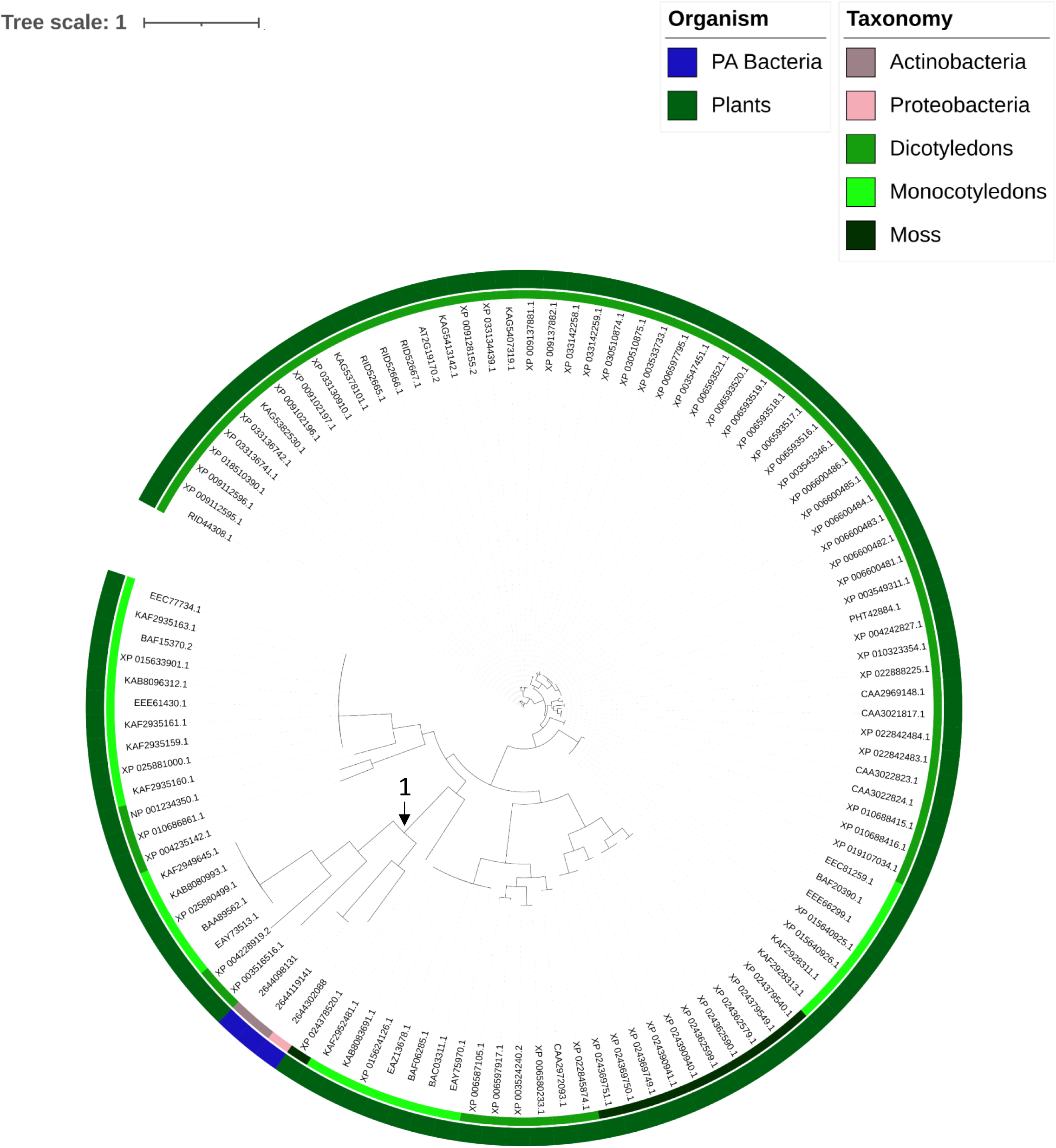
Example of HGT from plants to PA bacteria. A phylogenetic tree that presents homologs of the SLP3 (AT2G19170) gene. The bootstrap value of the clade that is shared by plants and their bacteria is 1 (marked with an arrow).

**Supplemental Figure 2.**
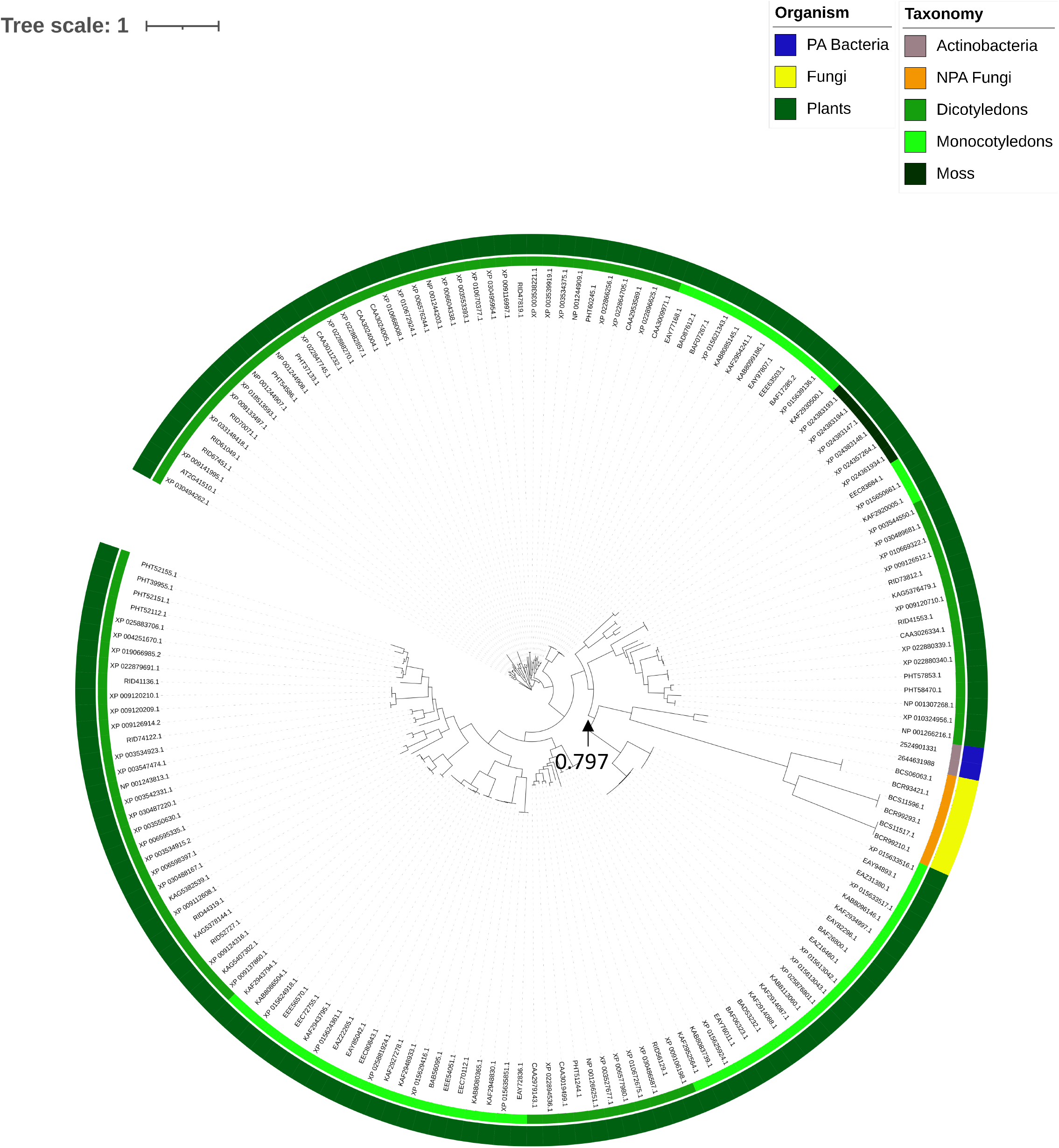
Example of HGT from plants to PA bacteria. A phylogenetic tree that presents homologs of the CKX1 gene (AT2G41510). The bootstrap value of the clade that is shared by plants and their bacteria is 0.797 (marked with an arrow).

**Supplemental Figure 3.**
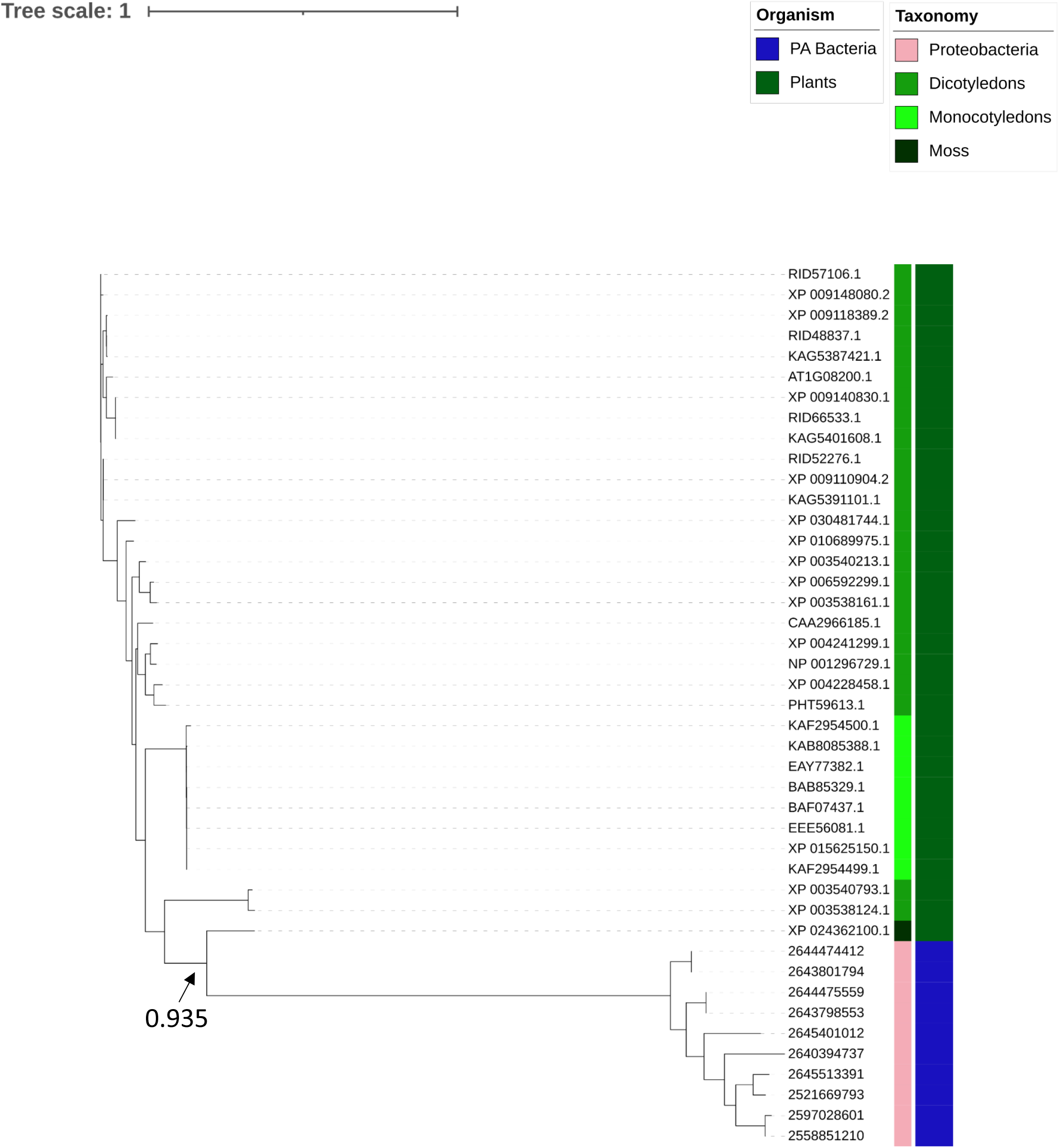
Example of HGT from plants to PA bacteria. A phylogenetic tree that presents homologs of the AXS2 (AT1G08200) gene. The bootstrap value of the clade that is shared by plants and their bacteria is 0.935 (marked with an arrow).

**Supplemental Figure 4.**
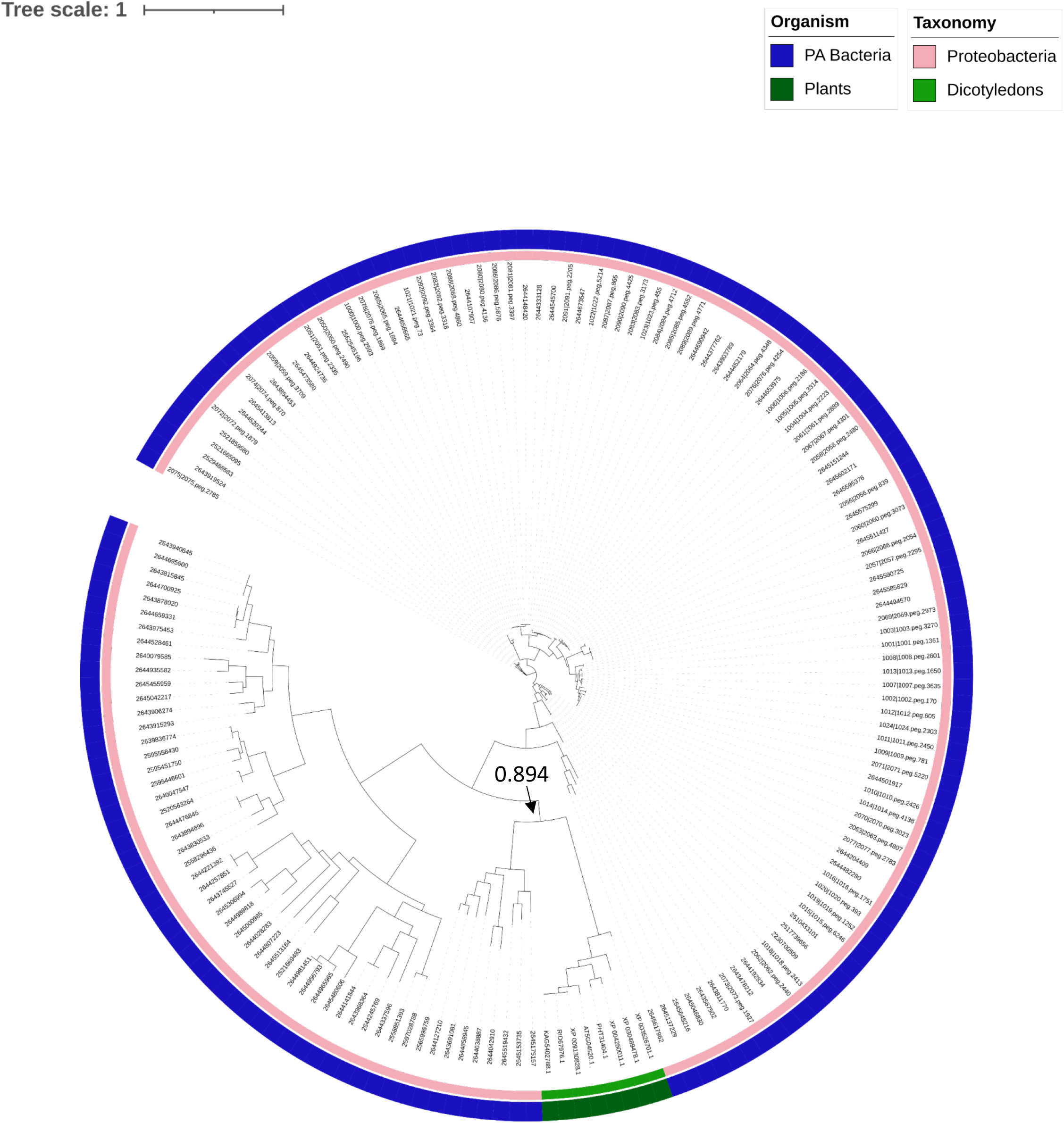
Example of HGT from PA bacteria to plants. A phylogenetic tree that presents homologs of the AT5G04520 gene. The bootstrap value of the clade that is shared by plants and their bacteria is 0.894 (marked with an arrow).

**Supplemental Figure 5.**
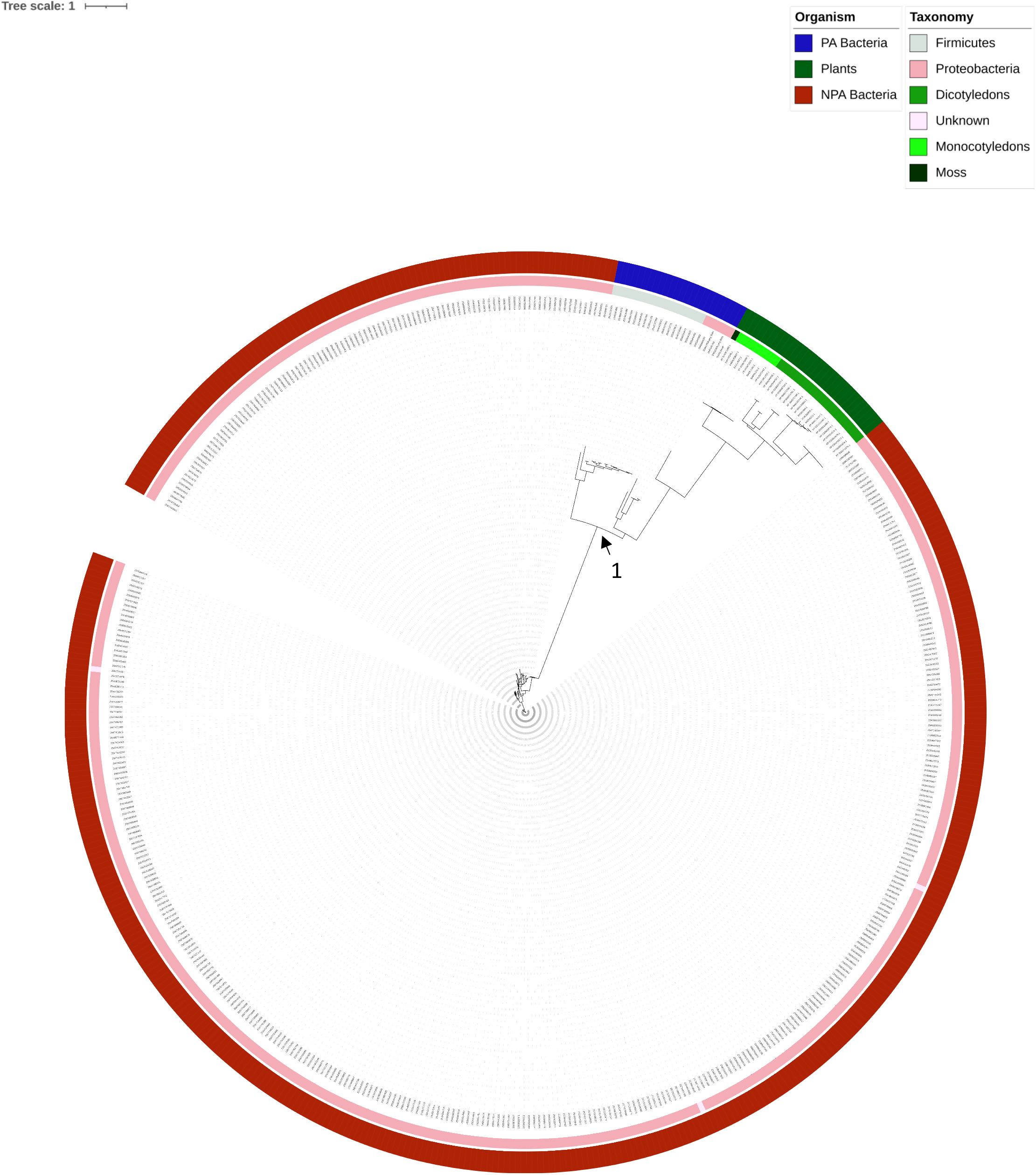
Example of HGT from PA bacteria to plants. A phylogenetic tree that presents homologs of the AT3G52905 gene. The bootstrap value of the clade that is shared by plants and their bacteria is 1 (marked with an arrow).

**Supplemental Figure 6.**
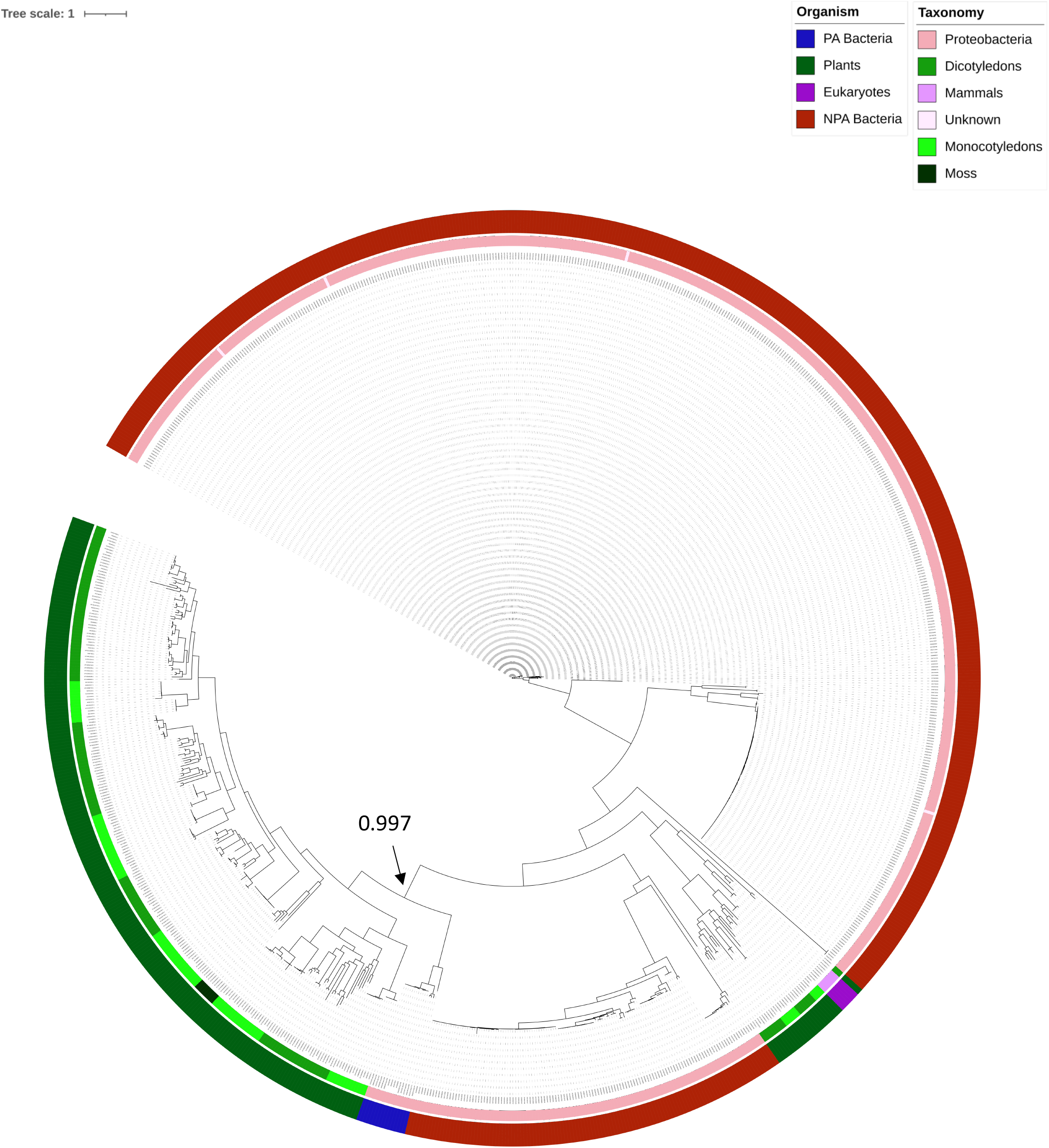
Example of HGT from PA bacteria to plants. A phylogenetic tree that presents homologs of the YUC8 (AT4G28720) gene. The bootstrap value of the clade that is shared by plants and their bacteria is 0.997 (marked with an arrow).

**Supplemental Figure 7.**
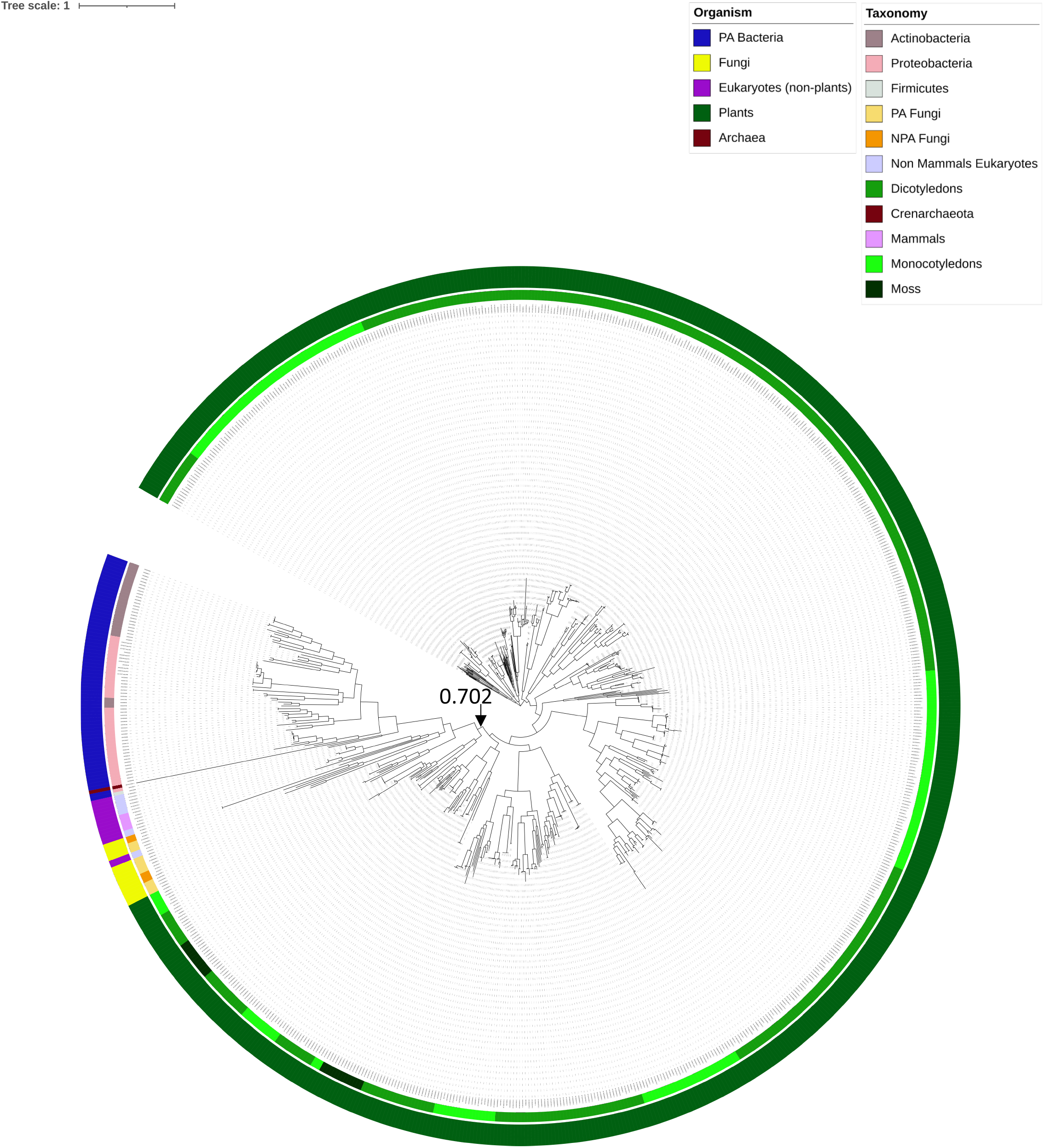
Example of HGT between eukaryotes and PA bacteria. A phylogenetic tree that presents homologs of the TGG4 (AT1G47600) gene. The bootstrap value of the clade that is shared by plants, their bacteria, and other eukaryotic organisms is 0.702 (marked with an arrow).

**Supplemental Figure 8.**
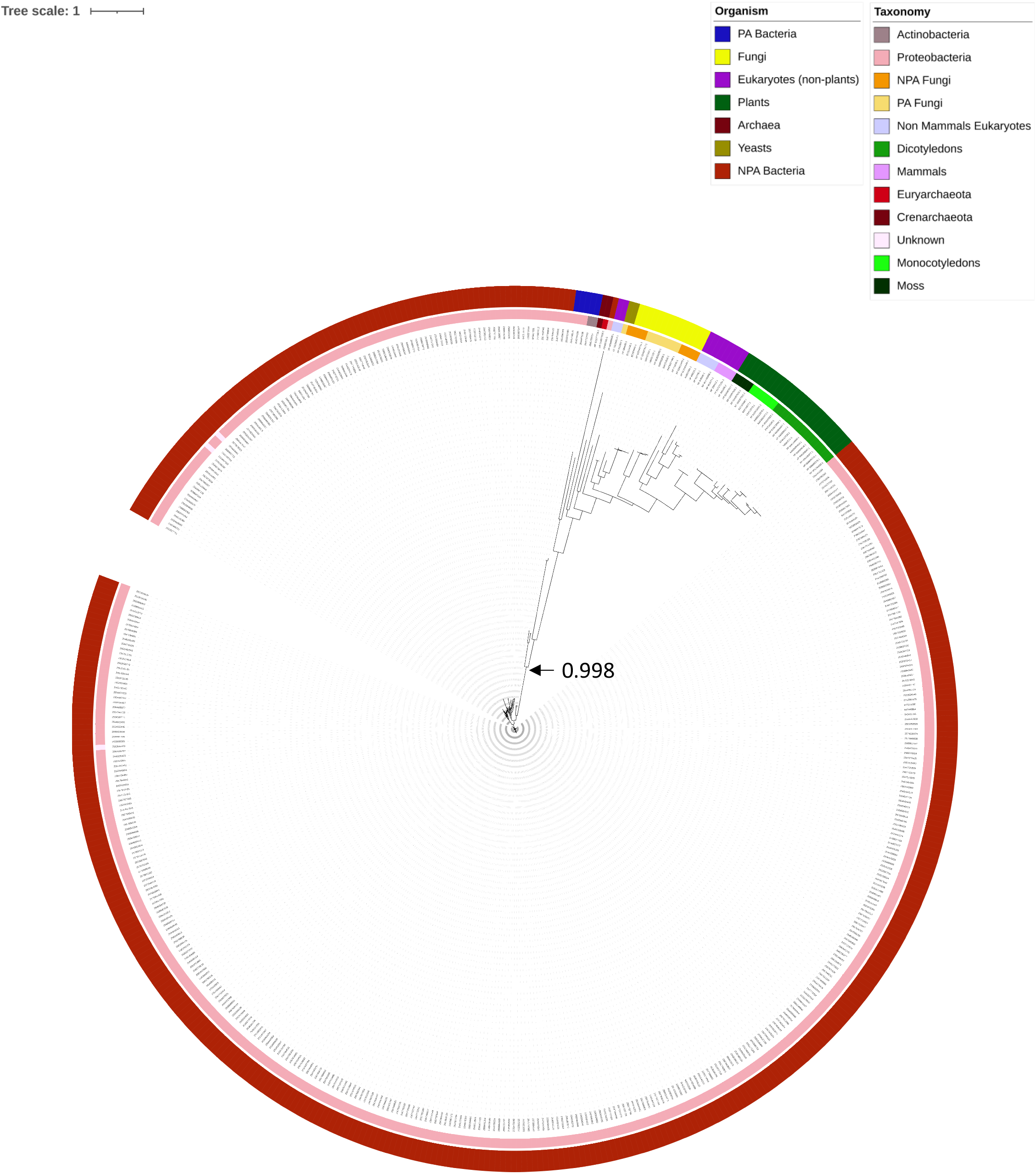
Example of HGT between eukaryotes and PA bacteria. A phylogenetic tree that presents homologs of the AT4G13720 gene. The bootstrap value of the clade that is shared by plants, their bacteria, and other eukaryotic organisms is 0.998 (marked with an arrow).

**Supplemental Figure 9.**
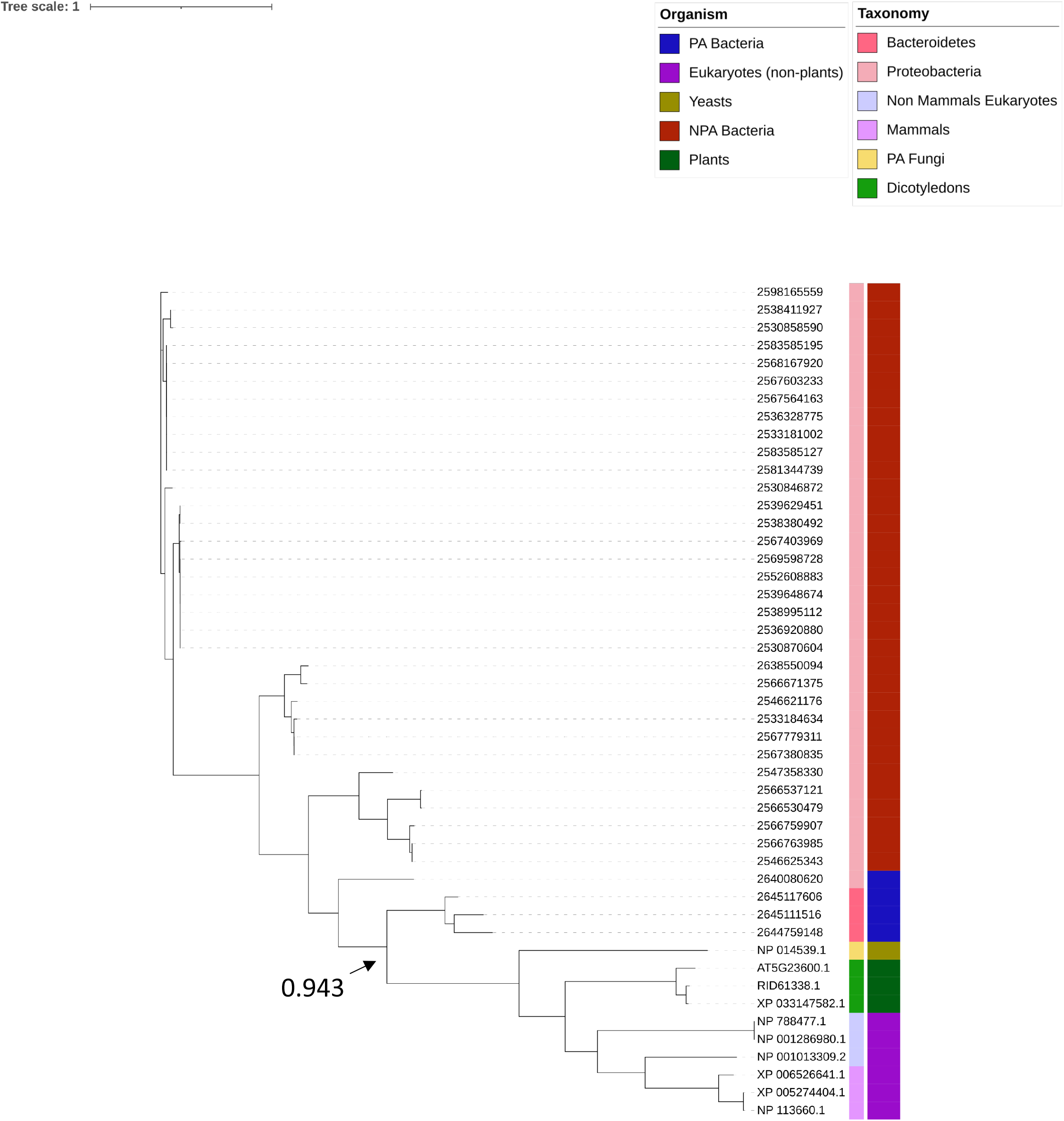
Example of HGT between eukaryotes and PA bacteria. A phylogenetic tree that presents homologs of the AT5G23600 gene. The bootstrap value of the clade that is shared by plants, their bacteria, and other eukaryotic organisms is 0.943 (marked with an arrow).

**Supplemental Figure 10.**
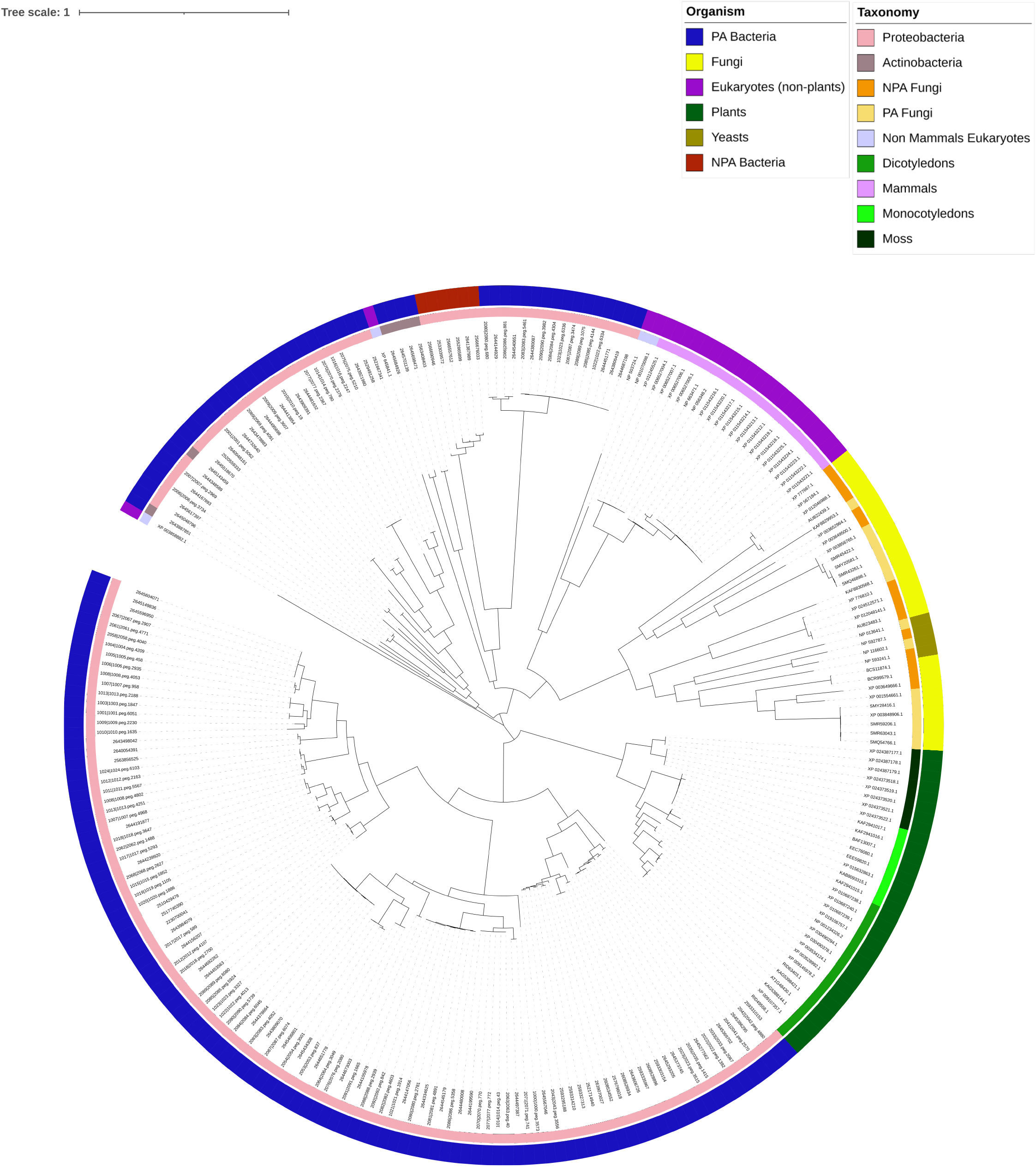
Example of unclear phylogenetic pattern. A phylogenetic tree that presents homologs of the AT1G48430 gene.

**Supplemental Figure 11.**
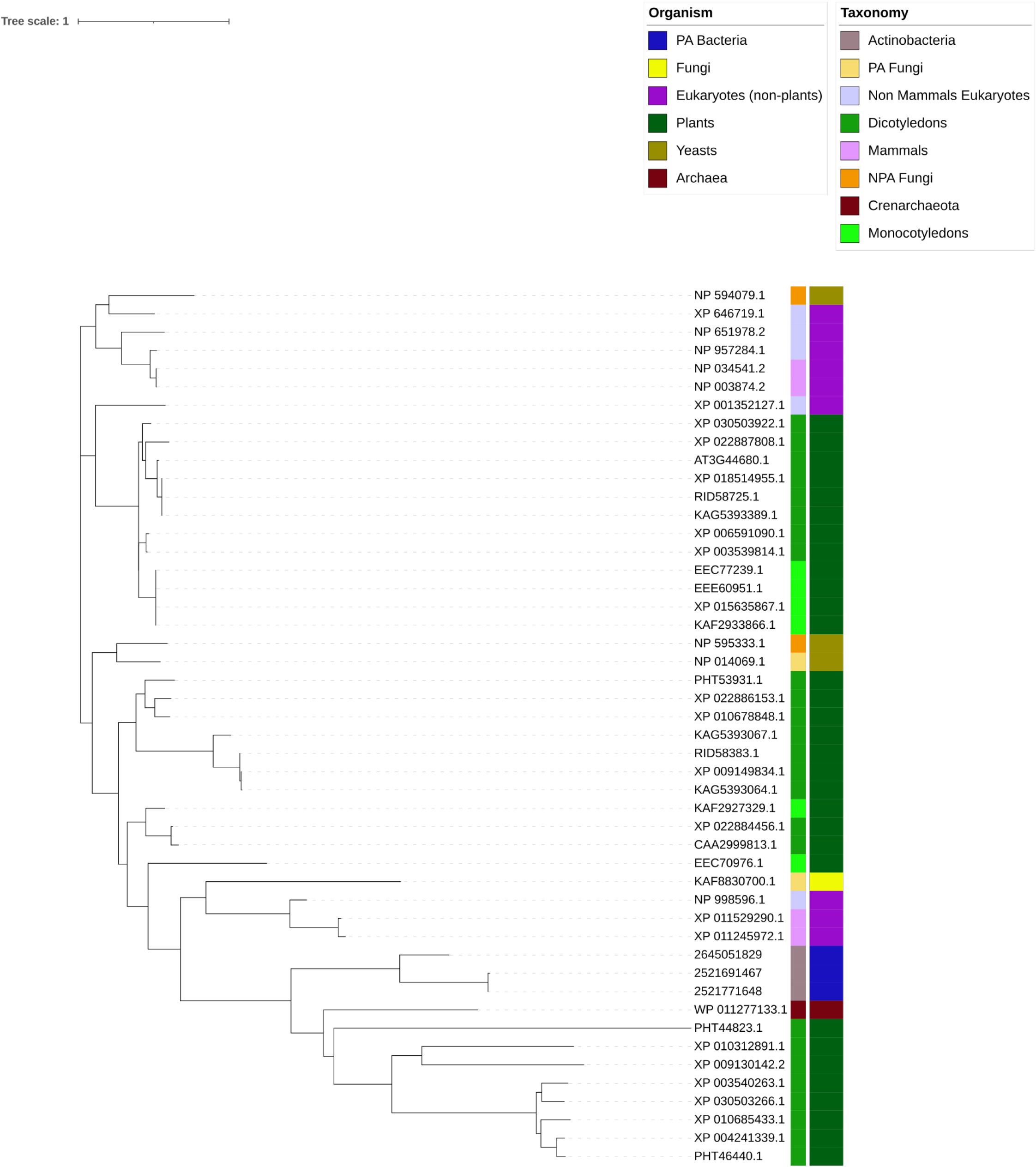
Example of unclear phylogenetic pattern. A phylogenetic tree that presents homologs of the HDA9 (AT3G44680) gene

**Supplemental Figure 12.**
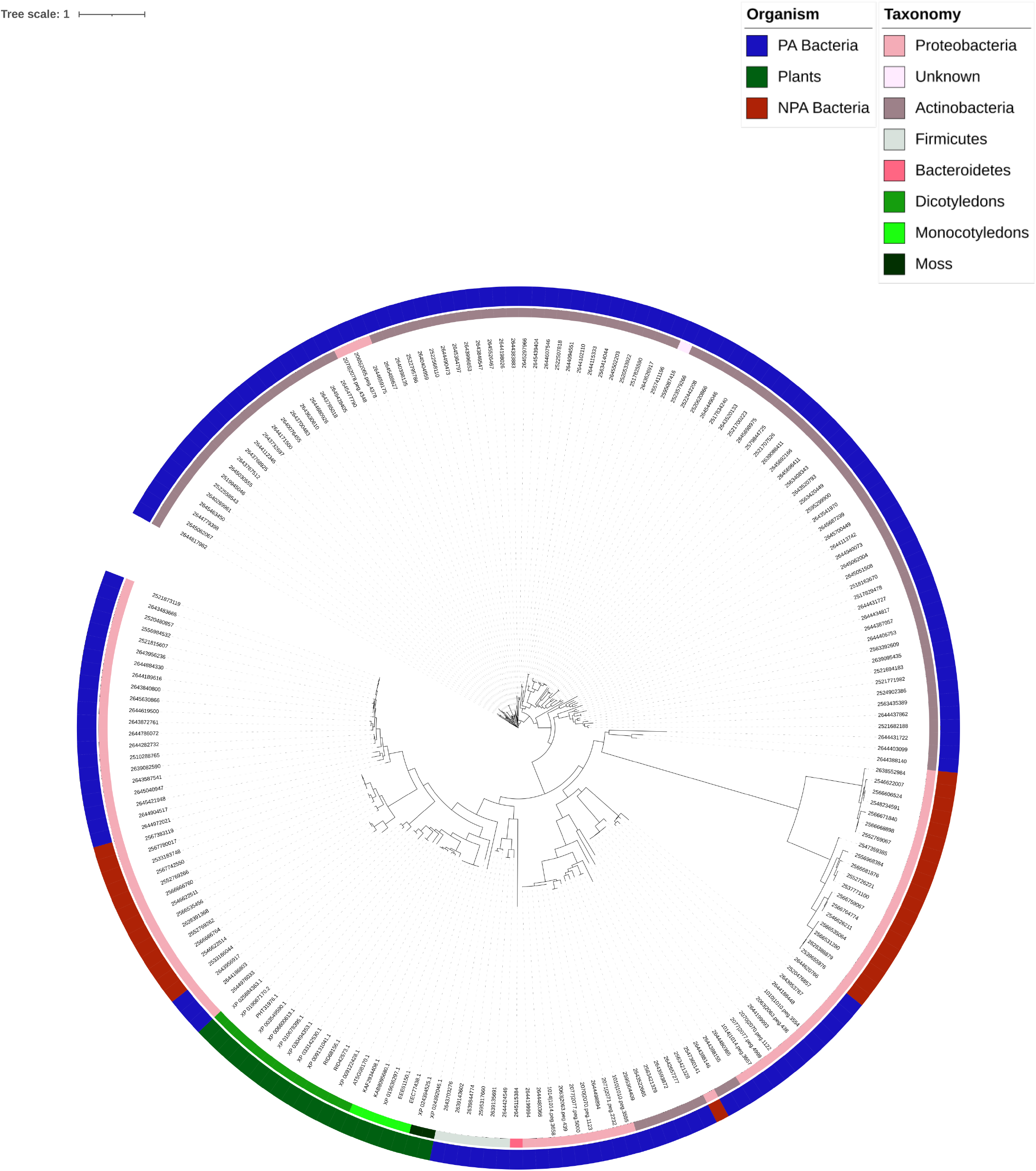
Example of unclear phylogenetic pattern. A phylogenetic tree that presents homologs of the ATKRS-1 (AT5G08170) gene.

**Supplemental Figure 13.**
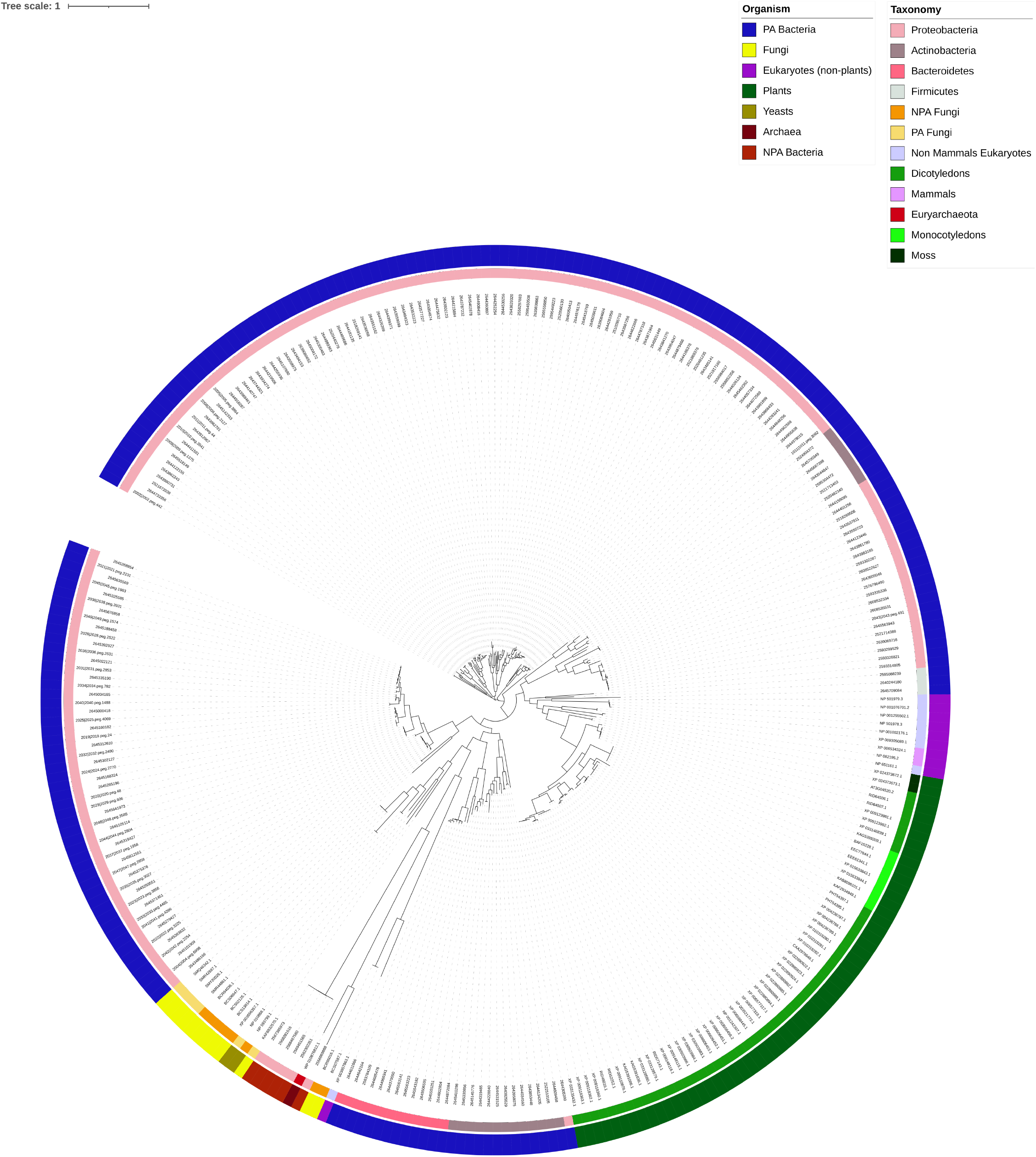
Example of no HGT detection. A phylogenetic tree that presents homologs of the THA2 (AT3G04520) gene.

**Supplemental Figure 14.**
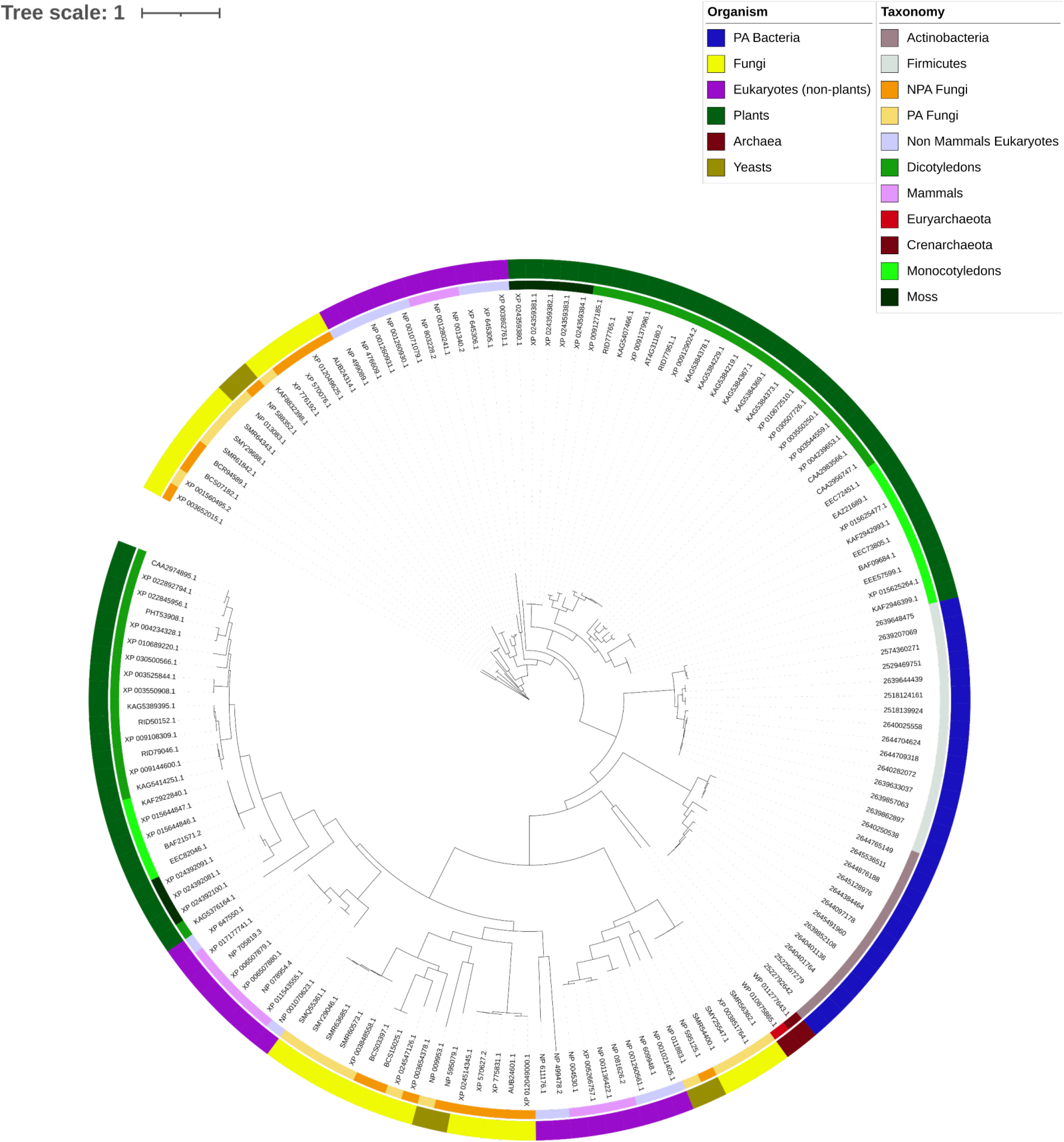
Example of no HGT detection. A phylogenetic tree that presents homologs of the IBI1 (AT4G31180) gene.

**Supplemental Figure 15.**
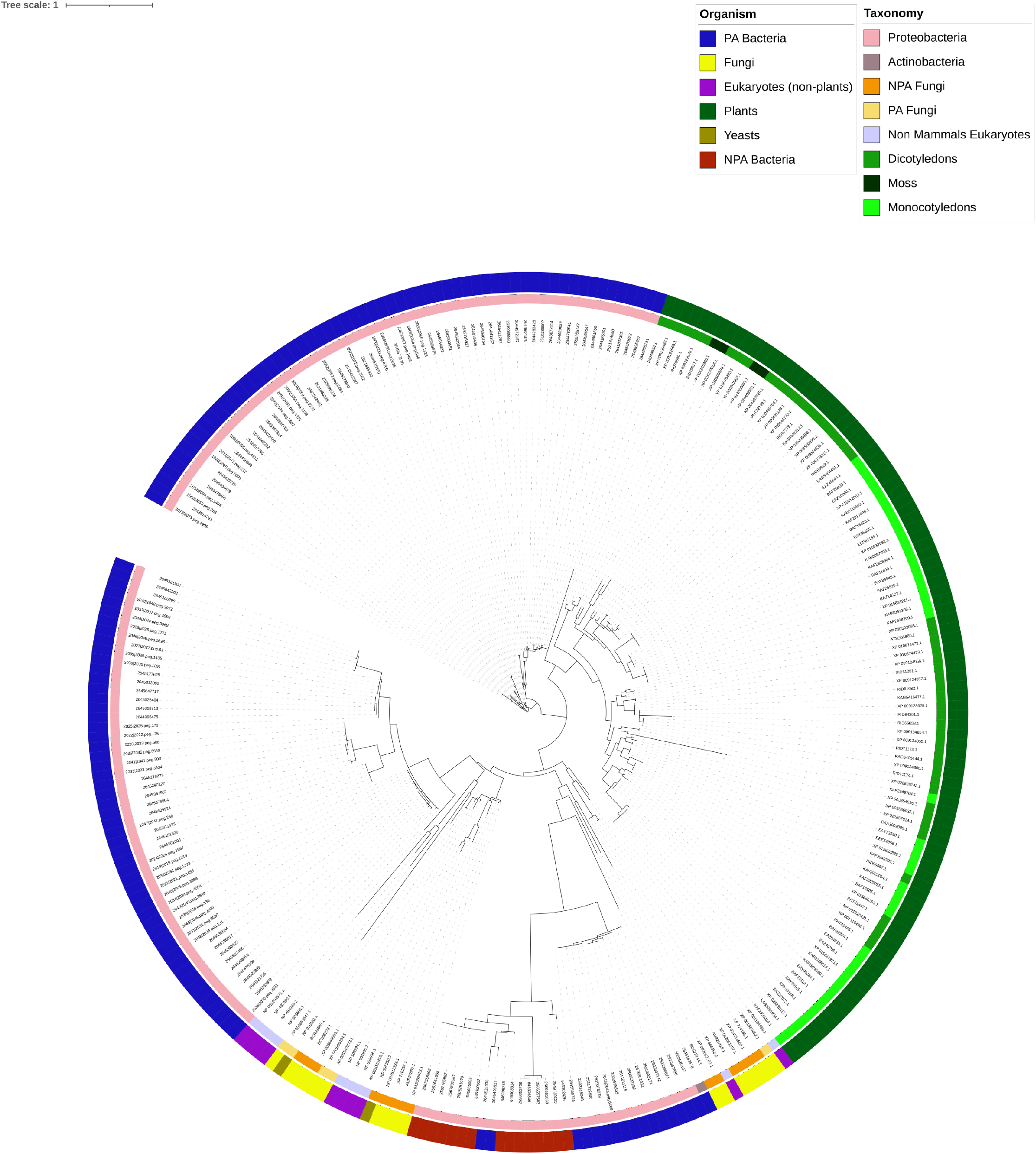
Example of no HGT detection. A phylogenetic tree that presents homologs of RCI2A (AT3G05880) gene.

**Supplemental Figure 16.**
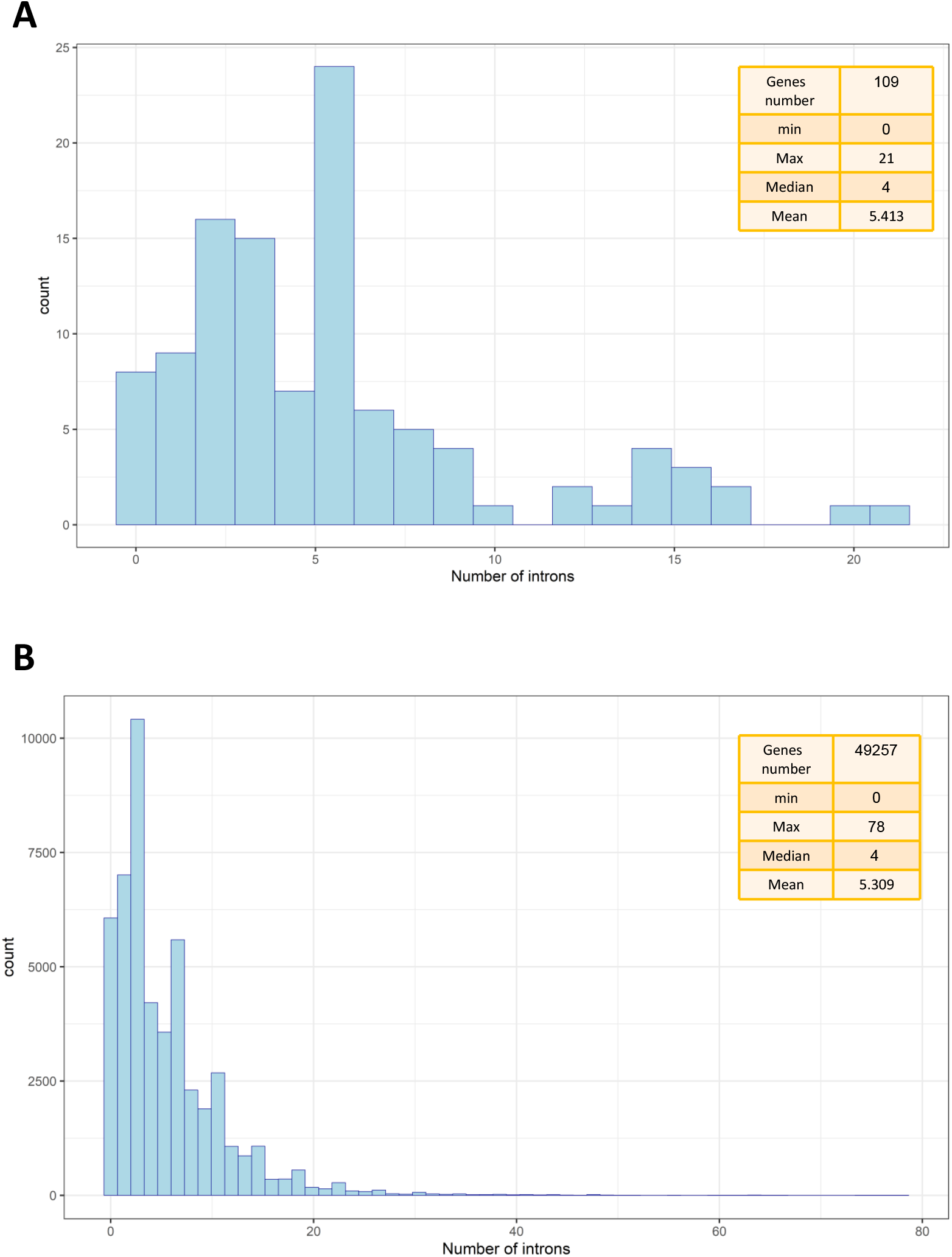
A. Number of introns of the genes belonging to the group “HGT from PA bacteria to plants”. B. Number of introns of all Arabidopsis.

**Supplemental Figure 17.**
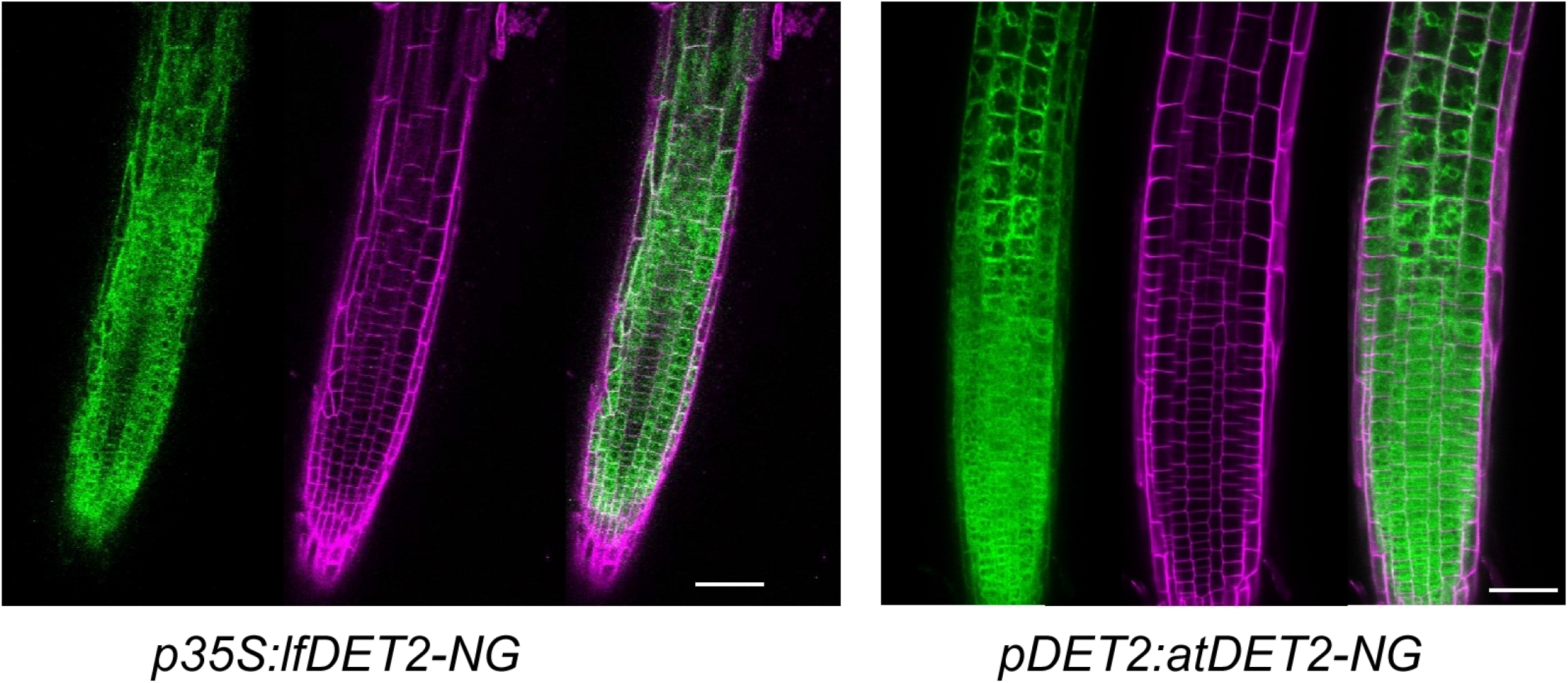
Confocal images of a roots expressing *lfDET2-NG* in the *det2* background (left) and *atDET2-NG* in WT background (right). NG is shown in green and propidium iodide (PI) that marks cell borders in magenta.

